# Diffraction-Limited Molecular Cluster Quantification with Bayesian Nonparametrics

**DOI:** 10.1101/2020.09.28.317057

**Authors:** J. Shepard Bryan, Ioannis Sgouralis, Steve Pressé

## Abstract

Life’s fundamental processes involve multiple molecules operating in close proximity within cells. To probe the composition and kinetics of molecular clusters confined within small (diffraction-limited) regions, experiments often report on the total fluorescence intensity simultaneously emitted from labeled molecules confined to such regions. Methods exist to enumerate total fluorophore numbers (e.g., step counting by photobleaching). However, methods aimed at step counting by photobleaching cannot treat photophysical dynamics in counting nor learn their associated kinetic rates. Here we propose a method to simultaneously enumerate fluorophores and determine their individual photophysical state trajectories. As the number of active (fluorescent) molecules at any given time is unknown, we rely on Bayesian nonparametrics and use specialized Monte Carlo algorithms to derive our estimates. Our formulation is benchmarked on synthetic and real data sets. While our focus here is on photophysical dynamics (in which labels transition between active and inactive states), such dynamics can also serve as a proxy for other types of dynamics such as assembly and disassembly kinetics of clusters. Similarly, while we focus on the case where all labels are initially fluorescent, other regimes, more appropriate to photoactivated localization microscopy, where fluorophores are instantiated in a non-fluorescent state, fall within the scope of the framework. As such, we provide a complete and versatile framework for the interpretation of complex time traces arising from the simultaneous activity of up to 100 fluorophores.

## 1 Introduction

Fluorescently labeled molecules, such as labeled proteins, are often used to create contrast between a cell’s background and the labeled molecular species of interest [1, 2, 3]. As biological processes unfold within cellular environments, these labeled molecular species may aggregate into clusters giving rise to the appearance of bright spots in fluorescence microscopy [4, 5, 6, 7]. Assessing the composition of these clusters on the basis of the spot’s variable brightness is a key step toward unraveling the role of molecular clusters [4, 5, 8, 9, 10, 6, 7].

Directly enumerating fluorophores and tracking their photophysical dynamics by discriminating between them on the basis of their physical location [11] is often impossible as typically an entire bright spot lies below the diffraction limit [2, 3]. Furthermore, fluorescence ruler methods, which enumerate fluorophores across time by comparing the brightness of a region of interest (ROI) to the brightness of a known calibration standard, are unreliable when the number of fluorophores is large on account of the inherent uncertainty introduced by photon shot noise which increases with growing fluorophore numbers [12, 13]. Other sources of uncertainty, beyond shot noise, include camera or detector noise and the rapid rise and fall of fluorescence intensity of the spot [4, 5]. The latter can arise on account of photophysical activity of the individual fluorophore labels as they cycle between fluorescently emitting or active (i.e., bright) and non-emitting or inactive (i.e., dark or photobleached) states [14] or it can arise due to assembly and disassembly of a cluster as individual constituents bind and unbind. For the purposes of this manuscript we will focus on brightness steps as being caused exclusively by photophysical dynamics and postpone further mention of (dis)assembly to the Discussion. A cartoon depicting this process is shown in figure 1.

**Figure 1:**
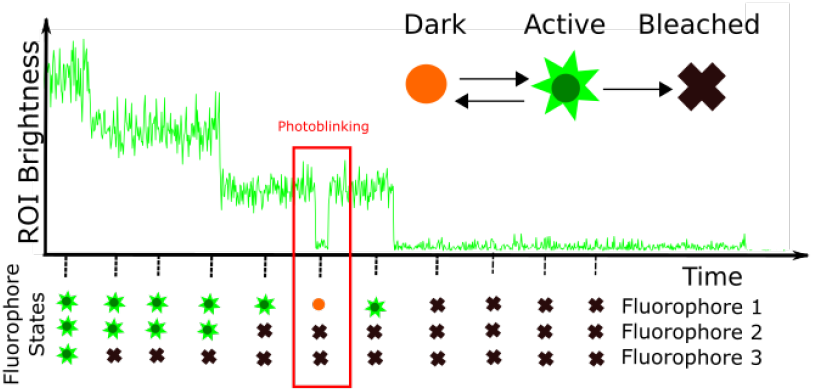
Summary of the problem. Each fluorophore attains one of three types of states: dark, active (bright), or photobleached. Only fluorophores in the active state emit photons. Each fluorophore transitions between these states as indicated by the arrows. The brightness over time of an ROI reflects the states of all fluorophores within the ROI. Our goal is to estimate the number of fluorophores in each ROI, as well as the photo-states of the fluorophores at each time level.

Traditionally, *Photobleaching step analysis* (PBSA) methods were developed to enumerate the number of fluorophores within a spot [12, 13, 15, 16, 17, 18, 19, 20, 21, 22, 23, 24, 25]. Such methods proceed in many ways; for example, by exploiting hidden Markov models [23, 25], data filtering to identify steps [17], statistical measures to identify expected violations of statistics characteristic of steps [21, 18, 20, 19], or neural nets [22]. In a recently submitted manuscript, additional ways of automating and improving upon PBSA methods listed above have also been explored [26]. Yet all PBSA are limited to clusters where fluorophores irreversibly inactivate one at a time until they are all photobleached giving rise to step-like transitions between brightness levels [27, 15, 2, 28, 29, 30, 31, 32, 33, 34, 35, 36, 37, 38, 39].

Our goal is to present a general framework that can simultaneously count and determine the photo-trajectories of fluorophores within a diffraction-limited ROI while taking into account photophysical artifacts such as blinking. To achieve this: 1) we exploit a realistic generative model that includes accurate photophysics, learns photophysical parameters, and can treat detailed camera models; 2) we relax the requirement that fluorophores all initialize from the bright state; and 3) we provide full Bayesian analysis providing not only point estimates, but also uncertainties over all unknown parameters. We show that the novel aspects of our method allow us to count upwards of 100 fluorophores in a single ROI. By virtue of the generality of our framework, we have the ability to treat other camera models or re-interpret the brightness step transitions as cluster assembly and disassembly kinetics. To illustrate our method, we use data in which a spot is illuminated with bright light and whose active fluorophores undergo photophysical transitions between bright and dark states before eventually photobleaching. As the number of fluorophores at any given time is unknown, we exploit tools within Bayesian nonparametrics [40, 41], in particular the Beta-Bernoulli process (BBP) [42, 43] never previously exploited in assessing the kinetics and composition of molecular clusters.

## 2 Methods

Here we set up: 1) the forward model, i.e., a model describing the stochastic dynamics of a large collection of molecules as well as other contributions to the data; and 2) discuss the inference strategy required to learn the number of fluorophores and their photophysical trajectories from the data. In particular, we show how we estimate the state (for simplicity we refer to photo-states as states) of each fluorophore at each time, the transition probabilities between fluorophore states, the probability that a fluorophore starts bright, the fluorophore mean brightness, and the background mean brightness. As with all methods within the Bayesian paradigm, whether parametric or nonparametric, we provide not only a point estimate for the maximum *a posteriori* (MAP) value of each variable, but also achieve full posterior inference with credible intervals.

### 2.1 Forward model

The forward model describes how the data are generated. We start with *R* diffraction-limited ROIs, indexed by *r* = 1,…, *R*. Each ROI has *K_r_* fluorophores indexed by *m* = 1,…, *K_r_*. We record the brightness (measured in ADUs) of each ROI for *N* successive time levels, indexed by *n* = 1,…, *N*. The brightness of the ROI *r*, at time level n is denoted 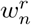 and is conditioned on the states of the fluorophores within the ROI at that time. The state of the *k*th fluorophore in ROI *r* at time level *n* is labeled 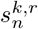. For simplicity, at each time level, we let 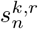 be in one of three states: dark, *σ_D_*, active, *σ_A_*, and photobleached, *σ_B_*. We tackle the obstacle of multiple bright states in the supplement (supplement 5.5).

At the first time level, each fluorophore in each ROI, starts either active or dark with probabilities given by ***π***_0_ which is an array with two elements: the probability of a fluorophore starting bright, *π*_0*A*_ and the probability of a fluorophore starting dark, *π*_0*D*_. At each following time level, *n*, the state of each fluorophore is conditioned on the previous state of the fluorophore according to ***π***, the transition probability matrix. Each element, *π_ij_*, of the matrix represents the probability that a fluorophore will be in state *σ_j_* given that it was previously in state *σ_i_* (supplement 5.2). These transitions include “dark to dark”, “dark to bright”, “bright to bright”, “bright to dark”, “bright to photobleached”, or “photobleached to photobleached” transitions (figure 1).

Our kinetic scheme is mapped here,

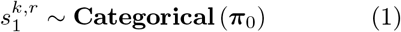

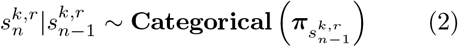

where 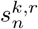 is the state of fluorophore *k* in ROI *r* at time level *n*, ~ means “is sampled from”, | means “given” or “conditioned on”, **Categorical** (*x*) means “the categorical distribution with probability mass *x*”, and 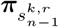 means “the row of **π** corresponding to the state of 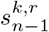”. The support for these categorical distributions is understood to be the set of possible states of the fluorophores, {*σ_D_, σ_A_, σ_B_*} meaning that for all *n, k*, and *r*, 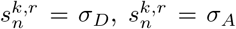, or 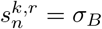.

At each time level, the fluorophores in each ROI give rise to the mean brightness of the ROI at the time level, 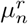. The mean brightness, 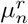 is the expected number of photons for the time level (calculated as the time step multiplied by mean photons emitted per unit time for the time level). We can decompose it into the sum of the mean background brightness of the ROI, 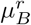, and the mean fluorophore brightness, *μ_A_*, multiplied by the number of active fluorophores in the ROI,

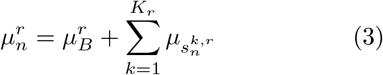

where 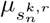 means “the brightness of the state corresponding to 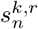”, as in, if 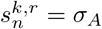 then 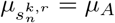 or if 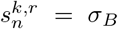 then 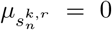. We note that 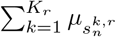 simply counts how many fluorophores are in the active state in the ROI at the time level of interest.

For data obtained with an EMCCD camera the brightness measured, 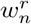, is conditioned on the mean brightness and the gain, *G*, through a gamma distribution [44]

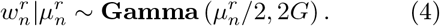

This model takes into account both shot and the readout noise [45]. Substituting Eq. (3) into Eq. (4) we find

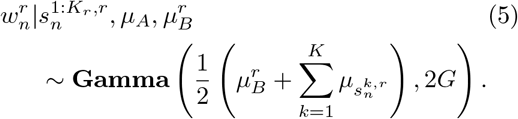

With this model, the mean expected readout is 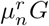 ADUs (units of camera readout) with a standard deviation of 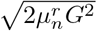 ADUs. Thus our model’s noise scales with the brightness with an excess noise factor of 2 that is characteristic of EMCCDs [46, 44]

This scheme, where many fluorophores give rise to a single measurement (the brightness at a time level), takes the form of a factorial hidden Markov model [47, 48].

### 2.2 Inverse formulation

We now develop the inverse formulation needed to estimate parameters from the data given a known number of ROIs, *R*, and associated time trace lengths, *N*. Following the Bayesian paradigm, we place prior distributions on all parameters whose posterior distribution we wish to determine. A graphical representation of our inverse model is shown in figure 2. Our choice of priors for transition rates and brightness parameters is straightforward and can be found in the SI (See supplemental section 5.1). However, our prior on the number of fluorophores is less straightforward as it requires a Bayesian nonparametric formulation that we outline below.

**Figure 2:**
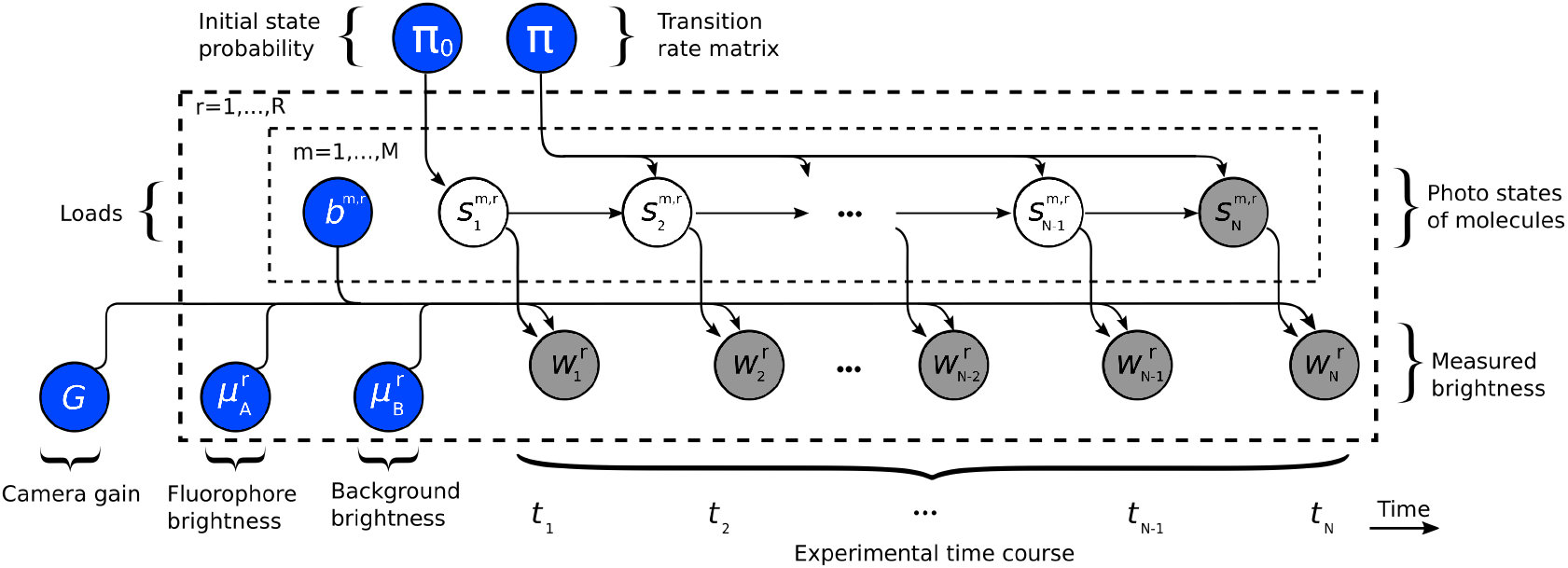
The graphical representation of the inverse model. Nodes (circles) in the graphical model represent random variables. Blue nodes are the random variables we infer. White nodes are auxiliary random variables (latent variables). Grey nodes are observations (data). The arrow between nodes indicates conditional dependence, meaning that if *x* is conditioned on *y* then we would draw an arrow going from node *y* to node *x*. The plates (dashed boxes) indicate that random variables within plates repeat over the index appearing at the top left of the plate. For example, the 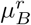 node is within the outer plate with index *r* implying an 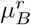 associated to each ROI (indexed *r*).

As we cannot set a prior on the number of fluorophores in each ROI, *K_r_*, we invoke Bayesian nonparametrics in our analysis. Briefly, we implement this using a nonparametric weak limit [49, 50]. That is, we assume an exceedingly large number of model fluorophores in the ROI, *K* ≫ *K_r_*, indexing each fluorophore with *k* = 1,…, *K*. We then assign each model fluorophore a load variable *b^k,r^*. If the load is on, *b^k,r^* = 1, we say that the fluorophore contributes to the ROI’s brightness. If the load is off, *b^k,r^* = 0, then the fluorophore is a virtual fluorophore which does not contribute to the brightness. Thus by summing the loads over all model fluorophores, we obtain the number of fluorophores located within the ROI. A load, *b^k,r^*, is a random variable sampled from the Bernoulli distribution with hyperparameter *γ*

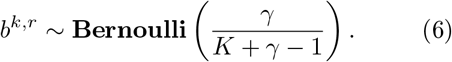

This probability mass is motivated by the Beta-Bernoulli process [42, 43] further discussed in supplement 5.4. In particular, as *K* becomes large, formally as *K* → ∞, the probability distribution converges to a distribution in which an infinite number of model fluorophores are considered [49, 50]. This choice of prior allows for inference independent on our choice for *K* provided a sufficiently large *K* (exceeding any reasonable number of fluorophores) is set; see supplement 5.10).

In analogy to equations (1)–(5), states are sampled just as we did in the forward model, except that each measurement, 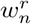, is now conditioned on the loads

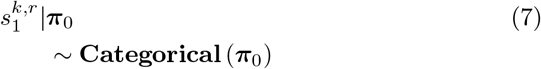

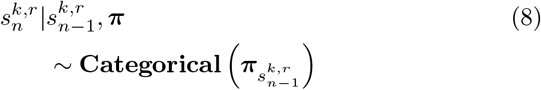

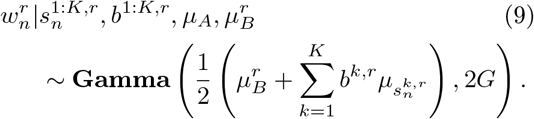

Here 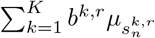 enumerates the number of fluorophores simultaneously active (i.e., loads in the active state in the ROI at time level *n*).

Lastly, if experiments are carried out long enough, all fluorophores eventually irreversibly photobleach. As such, we have knowledge of the final states of the fluorophores. Put differently, the fluorophore states at the last time level are fixed at

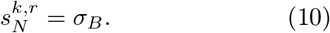

As such 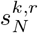 is shaded in grey in figure 2.

Together, these equations allow us to construct the high dimensional posterior over the collection of random variables (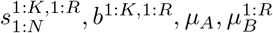, ***π***, and ***π***_0_)o This posterior does not assume an analytical form. As such, we employ the Markov chain Monte Carlo framework to computational sample parameters from this posterior [51, 52, 40, 53]. Briefly, our Gibbs sampler starts with an initial set of values for the parameters and attractively samples new values for each parameter one at a time while holding the others fixed (supplement 5.7).

## 3 Results

Here we demonstrate our method on simulated and experimental data for purposes of model validation. We show that we can accurately learn the number of fluorophores within in an ROI as well as the fluorophore photo-trajectories. We do so robustly even as the number of fluorophores approaches 100. In the supplement, we perform a more detailed robustness analysis on our method using simulated data (supplement 5.10). There, we test our method by varying the number of loads and the number of simulated fluorophores, and the fluorophore state model.

To validate our method on real data, we analyzed brightness traces where fluorophores undergo transitions between photophysical states as they eventually photobleach. This data uses Gattaquant DNA origami constructs with known number of fluorophore binding sites (such that ground truth be known on the total expected number of fluorophores) labelled with ATTO-647N fluorophores with known binding efficiency [26].

Traces with ATTO647N fluorophores examined by us and by others [54] show that ATTO647N has two bright states (see SI section 5.13 for plots of the data traces). We note that in the following sections, our model is supplemented to accommodate a second bright state for the fluorophores in the data we analyze. The expanded model is discussed in supplement 5.5.

### 3.1 Data acquisition

Data acquisition, provided by Hummert and Yserentant et al. [26], is briefly summarized here. ATTO647N labeled DNA oligomers were bound to DNA origami constructs. The DNA origami were imaged using a custom built Nikon Eclipse microscope with total internal reflection fluorescence (TIRF) illumination and a back illuminated EMCCD iXon Ultra 897 camera [26]. A log of Gaussian filter was used to select ROIs. Traces including artifacts such as diffusing fluorophores where excluded. For each ROI at each time level, we summed the brightness of every pixel within the ROI to get the total ROI’s brightness at each time level (i.e., the brightness time trace). We took time traces using two different types of of DNA origami constructs with 20 and 35 binding sites, respectively. For the 20 binding site origami, movies were taken for 1000 seconds at 50ms camera exposure (20000 frames) with a gain of 50. For the 35 binding site origami, movies were taken for 3000 seconds at 200ms camera exposure with an gain of 10 (15000 frames).

In order to analyze traces with more than 20 or 35 fluorophores, we also summed the brightness of every pixel involving multiple ROIs to get the total brightness arising from these combined ROIs at each time level. Because our camera model is a gamma distribution which is closed under addition, this procedure generates controlled traces with a ground truth containing known multiples of 20 or 35 fluorophores.

### 3.2 Results on simulated data

We evaluated our method with data simulated using the forward model put forward in Eqs. (1)–(5) with parameters chosen to mimic real data. We simulated 50 ROIs containing 14 fluorophores on average. The traces are 1000 *s* long with brightness 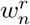 collected every 50 *ms*, so 20000 total frames. The exact number of fluorophores in each ROI is sampled from a binomial distribution to mimic 20 binding sites with 70% labeling efficiency. The gain used for the simulation was 50 [55]. The dimensionless background brightness parameters are 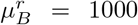. The fluorophores were simulated with two bright states with brightness given by *μ*_*A*1_ = 450 and *μ*_*A*2_ = 350 (plus one dark state and a photobleached state with brightness given by *μ_D_* = *μ_B_* = 0). These values were chosen to mimic the experimental data that we analyze in section 3.4. For example, the height and duration of a simulated photobleaching event qualitatively match those seen in the real data (see SI section 5.13).

Figure 3 shows the results for our analysis. The left panel shows the measured brightness versus time trace, superimposed with a sampled mean brightness over time, and the ground truth mean brightness over time. By mean brightness over time we mean the mean expected measurement at each time, 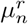, given the number of fluorophores in each state at that time level, the brightness of each state, and the camera gain. The mean brightness over time directly informs us on the photo-states of the fluorophores; see Eq. 3. Importantly, we capture all brightness drops due to blinking (i.e., photophysical dynamics) that cannot otherwise be obtained using existing PBSA methods that have built into them assumptions 1 and 2 discussed in the introduction.

**Figure 3:**
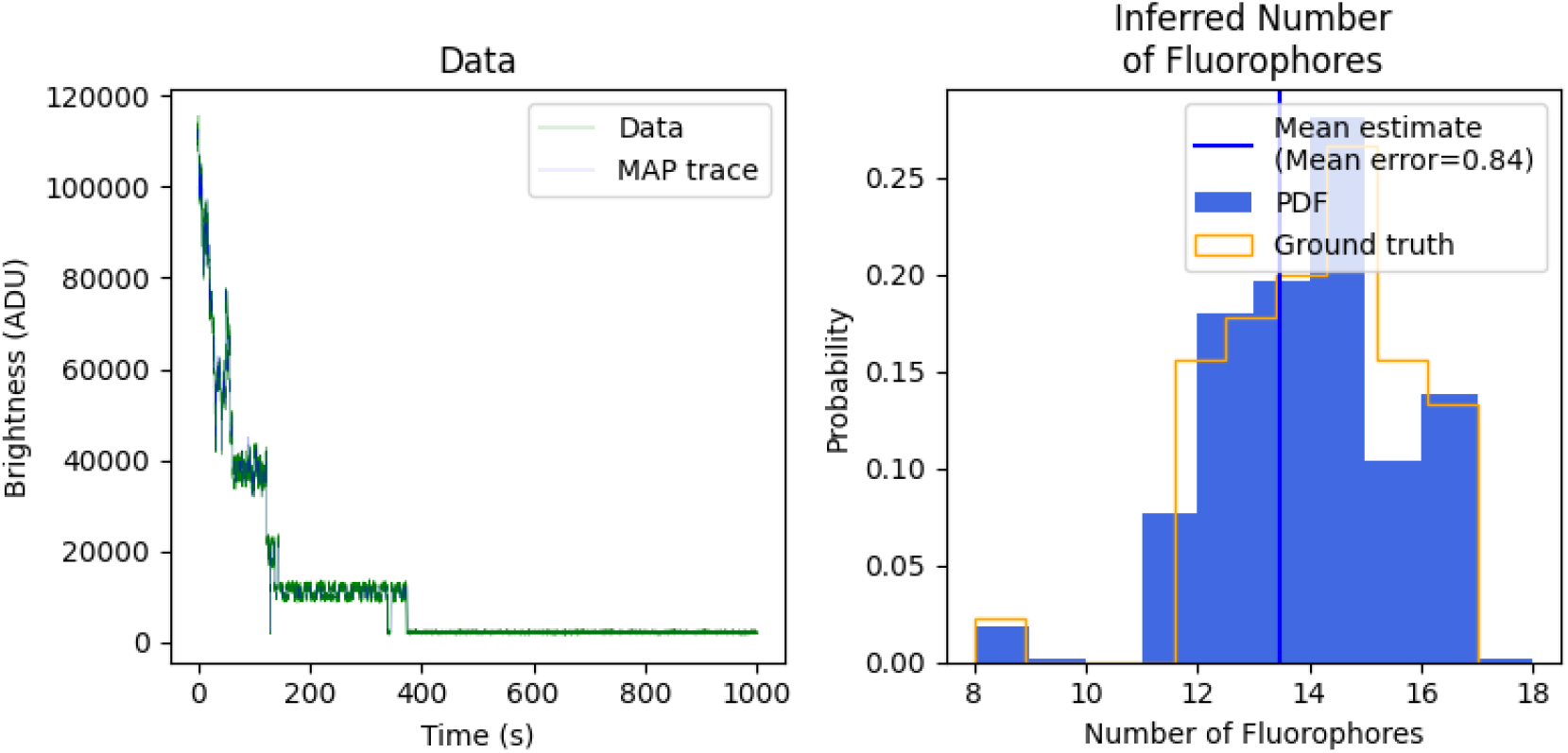
Inference on simulated data. We generated data using the forward model in section 2.1. We learn the number of fluorophores in each ROI and compare to ground truth (right panel) and associated photo-trajectories in each ROI (shown on left panel for one of many ROIs). In addition, we must also simultaneously and self-consistently learn all other associated parameters shown in figure 2.

On the right panel we show posterior over the number of fluorophores per ROI. That is, we find 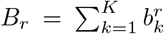 for each ROI and each sample of our posterior. We then histogram the *B_r_*’s for *r* = 1,.., *R*. In the limit that the number of ROIs is large, this should converge to the ground truth distribution of fluorophore numbers marginalized over the uncertainty associated with the number of fluorophores in a single ROI. We calculate the mean error of our method as the average difference between our estimate and the ground truth. Our sampled mean expected brightness trace matches well with the ground truth (within 1 fluorophore). Error analysis shows that roughly half of the samples were equal to the ground truth. No samples were more than 2 fluorophores off.

### 3.3 Comparison against other methods

Here we compare the results of our method to those obtained using the change point method of Tsekouras et al. [15], the two state model of Garry et al. [25], as well as a ruler method [12, 13]. We note that the Garry et al. method is equivalent to a two state implementation of our own method though they focus on state populations whereas we look at the state of each individual fluorophore. As such, we use our own method, but modified to include only one bright state, one photobleaching state, and no dark state, when comparing our method to the two state model. Our implementation of the ruler method is explained in SI section 5.9. We compare the methods on three different data sets: 1) data simulated using the same parameters as in the demonstration (section 3.2); 2) data simulated in which some fluorophores initiate in the dark state; and 3) data simulated with higher noise. Figure 4 shows the results of our comparison.

**Figure 4:**
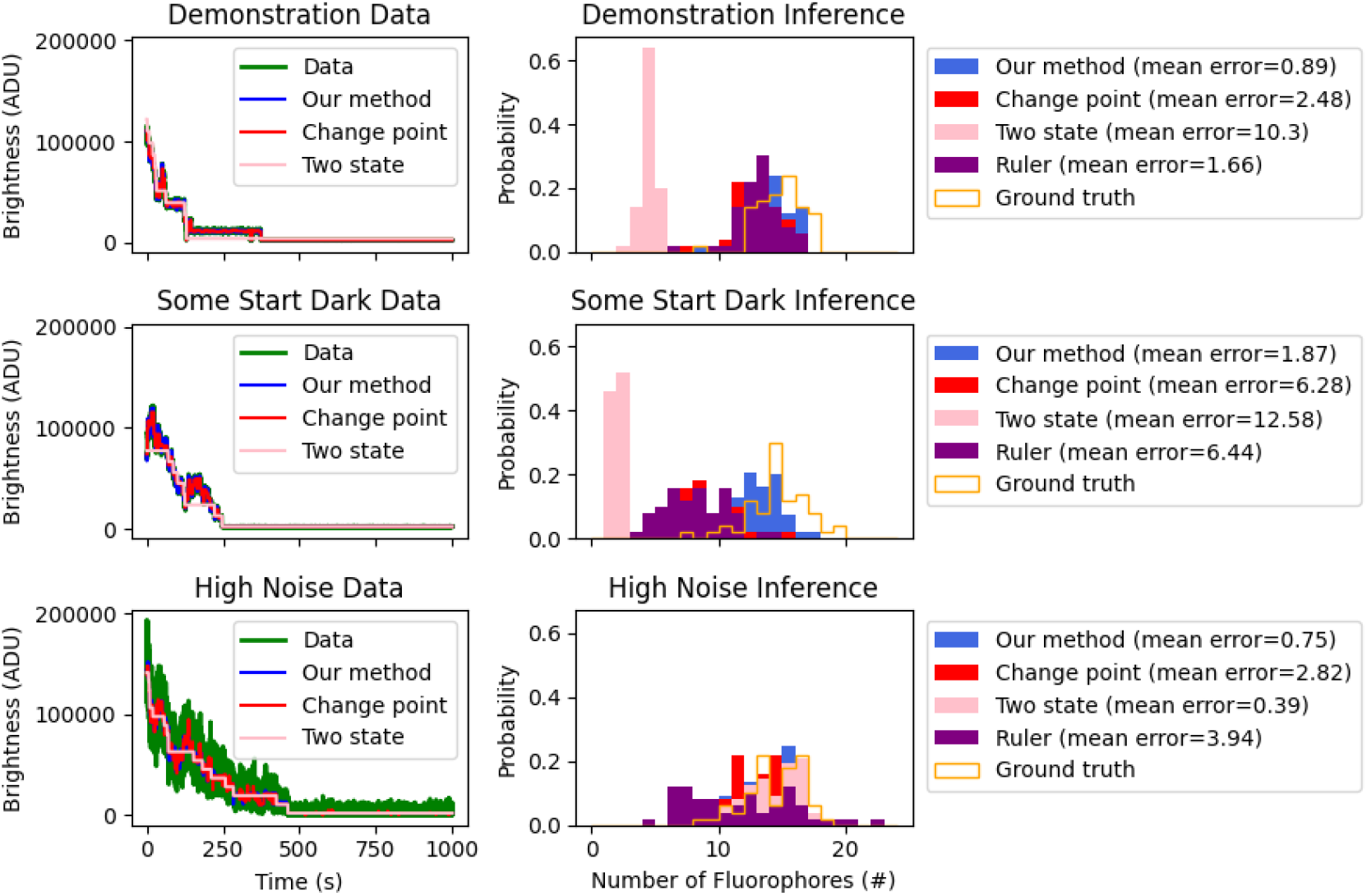
Comparison against other methods. Here we compare our method against a change point method, the ruler method, and the two state model. We compare all methods on three different data sets. On the top row we compare on simulated data using the base set of parameters. On the second row we compare on simulated data in which some fluorophores start dark. On the bottom we compare on simulated data with high noise. The left panel of each row shows the inferred phototrajectory for an ROI using our method, the change point method, and the two state model (the ruler method does not generate trajectories). The right panel shows the inferred distribution for the number of fluorophores for each of the different methods. In the legend we show the mean error of each method calculated as the average difference between the inferred number of fluorophores and the ground truth.

As seen in the top row of figure 4, all three methods, besides the two state model, do reasonably well (within 20% error) in inferring the number of fluorophores using the base set of parameters. The two state model underestimates the number of fluorophores due to the fact that it cannot account for blink events (see SI section 5.10.3). Note that the mean error (the average difference between the estimated number of fluorophores in an ROI and the ground truth number of fluorophores used in the simulation) was smallest for our method. That all four methods do well is expected because the data is clean and the steps are easy to see by eye and therefore all three methods should do well at identifying brightness levels and inferring the number of fluorophores.

Next we look at simulated data in which some (40%) of the fluorophores start in the dark state. The second row of figure 4 shows the results. Here, the two state model, the ruler method, and the change point method underestimate the number of fluorophores by over 40% because they do not allow for fluorophores to initiate in a dark. Our method, which allows fluorophores to initiate in such a state, learns the number of fluorophores with less than two fluorophores mean error.

The last row of figure 4 shows results on data simulated with higher noise. The higher noise level was achieved by decreasing the brightnesses, *μ_A_* and *μ_B_*. This physically represents lowering the intensity of the laser used to excite the fluorophores down to a level where shot noise dominates. We simultaneously raised the gain to keep the average brightness at the same level. Under these conditions, the two state model no longer underestimates the number of fluorophores as brightness drops arising from blinking events are within the variance of the noise. As such, the two state model becomes reasonable. Our model, which has four states including two bright states and a dark state, has negligibly greater mean error than the two state model in this experiment, due to slight overfitting from having two bright states with brightnesses very close to each other relative to the measurement noise. While the two state model was able to infer the number of fluorophores in the noisy data essentially as accurately as our method, it was unable to do so in the low noise limit (figure 4 top row). As such, our physically-inspired method with photophysical dynamics reveals itself to be most robust across a range of scenarios.

### 3.4 Results on experimental data

Results from experimental data are shown in the top left and bottom left of figure 5. Here we plot the inferred distribution for the number of fluorophores in an ROI against the ground truth distribution for the number of fluorophores. The ground truth distribution of fluorophores here is binomially distributed [48] assuming a 70% percent labeling efficiency. The 70% labeling efficiency was provided to us by the manufacturer.

**Figure 5:**
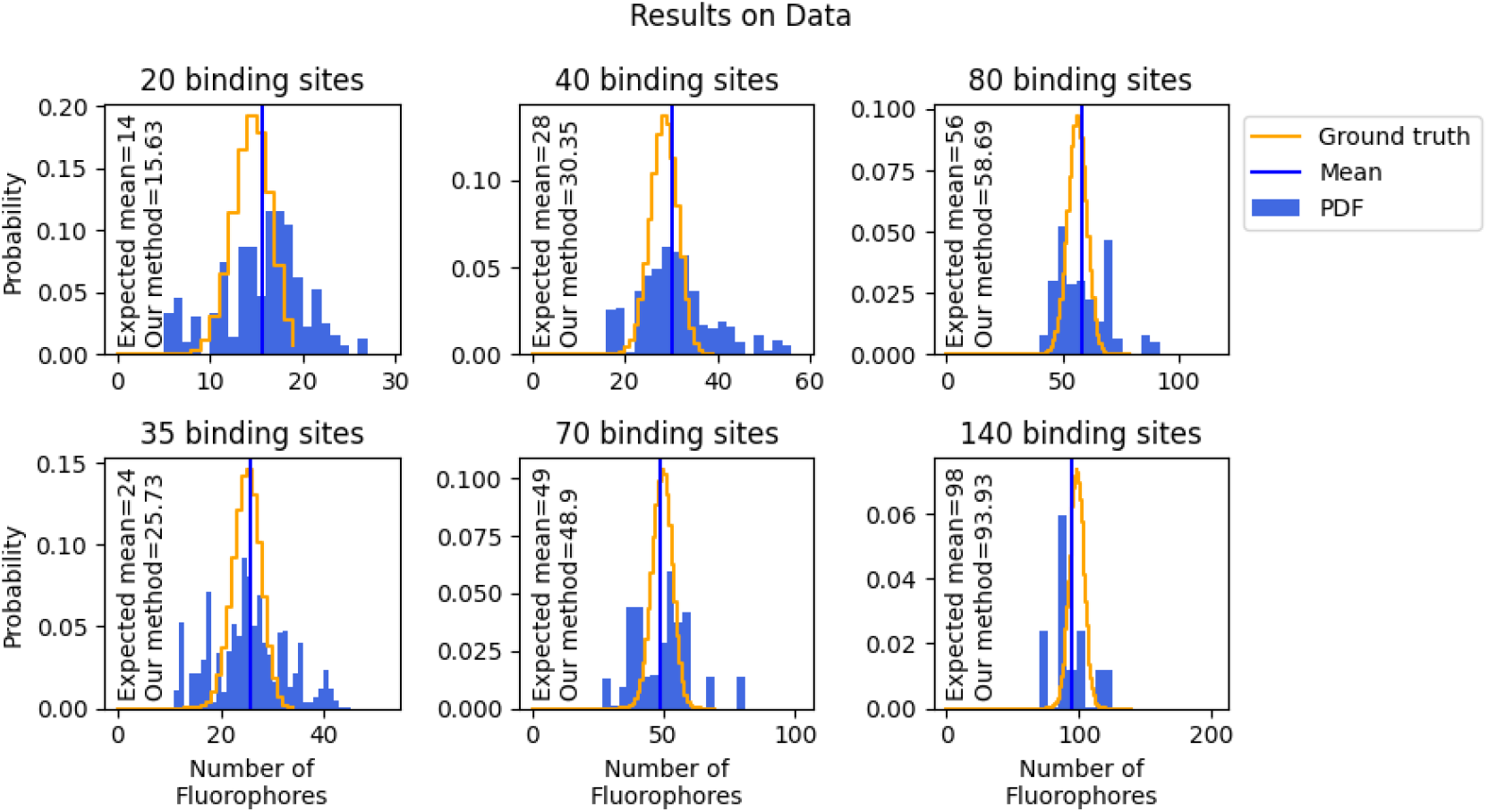
Inference on real data. Here we illustrate our method in enumerating fluorophores from real data. The top row analyzes data from experiments using DNA origami with 20 binding sites (we then combine data from different ROIs to generate data sets with a higher number of fluorophores). Similarly, the bottom row analyzes data from experiments using DNA origami with 35 binding sites (also combined to form ROIs with a larger number of fluorophores). Also plotted are the expected distribution of fluorophores (a binomial distribution), and a vertical line showing the mean expected number of fluorophores learned from our method.

We note that the width of the ground truth distribution for the number of fluorophores in the ROI arises due to labeling efficiency of the fluorophores, whereas the width in the distribution of the learned number of fluorophores arises from labeling efficiency as well as uncertainty in the inference. As such, we expect the distribution over the learned number of fluorophores to naturally be wider than the ground truth distribution for the number of fluorophores. For example, in the extreme case where we had 100% labeling efficiency, the ground truth distribution would have zero width, yet our method would still have a width due to uncertainty in the estimate. On the other hand, the mean estimated number of fluorophores in each ROI should be close to the ground truth and thus remains a reliable way by which to evaluate the accuracy of our method.

For our 20 binding site analysis, the predicted mean of the distribution for the number of fluorophores is only about 1.3 fluorophores higher than expected as can be seen in the top left panel of figure 5. This is likely due to overfitting sources of noise not accounted for in our model such as unbound fluorophores freely diffusing above the origami structure.

Given the agreement between ground truth and our method for 20 and 35 binding sites, we wanted to test how high we could count. In order to create controlled data sets with known ground truth, we combined the data from ROIs as discussed in section 3.1. For example, by summing together two ROIs with 20 or 35 binding sites, we could count fluorophores in ROIs with as many as 40 or 70 total binding sites (figure 5 middle column). By adding together four ROIs with 20 or 35 binding sites, we could generate new ROIs with as many as 80 or 140 fluorophores (figure 5 right column). For all four cases, the mean number of fluorophores per ROI learned from our an analysis closely matches (within 3 fluorophores) the ground truth of the expected mean.

## 4 Discussion

Learning the number of molecules located within a molecular cluster, while simultaneously and self-consistently determining the dynamics of the cluster’s constituent members, is a key step toward unraveling life’s processes occurring well below light’s diffraction limit [27, 15, 2, 28, 29, 30, 31, 32, 33, 34, 35, 36, 37, 38, 39, 12, 13, 15, 16, 17, 18, 19, 20, 21, 22, 23, 24, 56]. In order to do so, we introduced a Bayesian nonparametric framework that accurately models the photophysics, shot noise, and detector noise that gives rise to the data, along with sampling methods capable of exploring this high dimensional probability space. Our method was illustrated for as many as 100 fluorophores. We note that the ability to count such a high number of fluorophores is necessary for cellular applications as, for example, nuclear pore complexes are known to be made up from 32 monomers [6, 26], Rac1 can aggregate into clusters of 50-100 [7], and Pol-II can aggregate into clusters of a few hundred [4].

By operating within the Bayesian paradigm, we can propagate uncertainty arising from sources of error, such as photon shot noise and detector signal amplification, into the full distributions over fluorophore numbers and the transition probabilities we determined.

Now, if the counting of fluorophores in a cluster were the only goal and it could be assumed that all fluorophores were initially active, then we could ignore dynamics altogether and avoid learning transition probabilities (as well as trajectories). In this case, a collapsed state formulation (one that keeps track of the total population of decreasing numbers of fluorophores) can be used [15, 25, 26]. However, even then, existing methods for enumeration do not sample full Bayesian posteriors and counting would not be possible for cases where the majority of fluorophores are initially inactive such as in the case of photoactivation localization microscopy (PALM) [57, 58, 59, 11]. Indeed, moving forward, PALM and other superresolution experiments [4, 5] could provide exciting in vivo test beds for our method.

Furthermore, while we have chosen to focus on brightness traces recorded using an EMCCD camera, we could in principle modify our method to allow other detector models. This could be achieved trivially be modifying equation 31 to incorporate the noise model of the desired detector. Moving forward this would allow photobleaching enumeration on a variety of detectors including photomultiplier tubes [60] or sCMOS cameras [61].

The generality afforded by our method in learning dynamics, and thus learning the state of every constituent member of a cluster explicitly, does come at an added computational cost. The majority of the computational cost comes from the forward-backward filtering algorithm used to sample the states. The forward filter backwards sample algorithm (FFBS) runs with time complexity 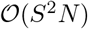 where *S* is the size of the the state space and *N* is the number of time levels. As we must run the FFBS over each load in each ROI, the total computation time to sample all the states scales like 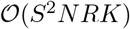 where *R* is the number of ROIs and *K* is the number of loads per ROI. Additionally, in order to facilitate proper mixing of the variables, we sample the states, two loads at a time in a joint state space of size *S^J^* where *J* is the number of loads we sample jointly (see SI section 5.7) which increases the size of the state space, but also decreases the number of times we have to run the FFBS per ROI (for example, sampling two loads at a time means we use FFBS half as many times). As such, overall, the time complexity of our algorithm scales as 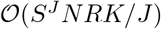. As the majority of the computational bottleneck is ascribed to sampling the states, we therefore sample the states of each ROI in parallel. Computational time can be improved by a factor of *R* if at each iteration of the Gibbs sampler, we sample the states for each ROI (which are independent from each other) in parallel. As the remaining parameters are sampled relatively quickly as compared to state sampling, we sample those globally at each iteration of the Gibbs sampler.

Finally, while we have focused on photophysical dynamics, it is possible to imagine learning the assembly and disassembly kinetics of a cluster. For example, using a two state model where the fluorophores transition between being cluster bound and unbound, our framework could be used to learn the state transition rates as well as the total number of fluorophores bound to the cluster at any given time. Learning such kinetics would be especially relevant to monitoring the formation of large transient protein assemblies relevant to cellular transcription [8, 9, 10]. What remains to be seen is how data could be analyzed if assembly and disassembly of molecules in a cluster are occurring while photophysics of labels on these molecules is simultaneously taking place. In this case, either stable fluorophores that remain in a bright state would need to be used or a difference in timescales between the assembly and disassembly kinetics and photophysical kinetics would need to be sufficiently large to be independently determined by a future analysis method.

## Author Contributions

SP and IS conceived of the project. JSB carried out the coding and development. SP oversaw all aspects of the project.

## Acknowledgements

We would like to thank Dirk-Peter Herten, Klaus Yserentant, and Johan Hummert for their collaboration and excellent data. S.P. acknowledge support from the NIH (grant no. R01GM134426 and R01GM130745) and NSF (Award no. 1719537).

## Data Availability

The data analyzed in this project was provided by Dirk-Peter Herten, Klaus Yserentant, and Johan Hummert [26]. Brightness traces from ROIs used in this manuscript can be found at http://statphysbio.physics.asu.edu/.

## Code Availability

Code can be found at http://statphysbio.physics.asu.edu/.

## 5 Supplemental Information

### 5.1 Choice of priors

Priors on initial state probabilities and priors on the rows of the transition probability matrix follow Dirichlet distributions [48, 62, 63, 64]

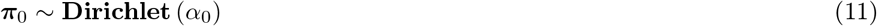

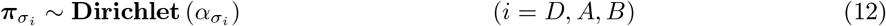

where *α*_0_ and *α_σ_i__* are hyperparameters (discussed further in supplement 5.7). The Dirichlet distributions are chosen for computational convenience alone as these are conjugate to the categorical distribution, Eq. (2); see supplement 5.3. The choice of prior becomes largely immaterial as the data set size increases.

We select gamma distributions as our priors on the mean fluorophore brightness and mean background brightness

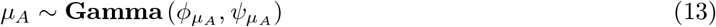

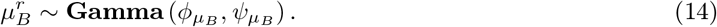

The gamma distribution choice reflects our knowledge that these variables are positive and the hyperparameters are calibrated in such a way as to broaden the range of values these brightnesses may adopt. In particular, by selecting small hyperparameters (*ϕ_μ_A__, ϕ_μ_B__* = 2), the prior gives non-negligible probability to a wide range of values for *μ_A_* and 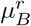; the choice for these scale parameters of the gamma distribution, *ψ_μ_A__* and *ψ_μ_B__*, is further discussed in supplement 5.7. Once more, the choice of hyperparameters become increasingly unimportant as more data are collected [40, 65].

### 5.2 Remark on transition probabilities

As shown in figure 1, fluorophores can transition from “dark to bright”, “bright to dark”, and “bright to photobleached” states as well as self transition from “dark to dark”, “bright to bright”, and “photobleached to photobleached” states. This is captured by the layout of ***π***, where rows correspond to the “old state” and columns correspond to the “new state”.

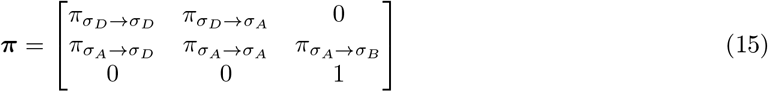

Here, *π_σ_B_→σ_B__* = 1 because there are no transitions out of the photobleached state. Similarly, ***π***_0_, the initial state probability, takes the form

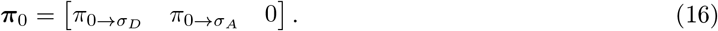

### 5.3 Conjugate prior on transition probabilities

The choice of Dirichlet prior on the rows of ***π*** combined with the likelihood give back a Dirichlet distribution [48].

*Proof:*

Using Bayes’s theorem, and dropping all terms that do not depend on ***π***_0_ and ***π***, the likelihood of our model is

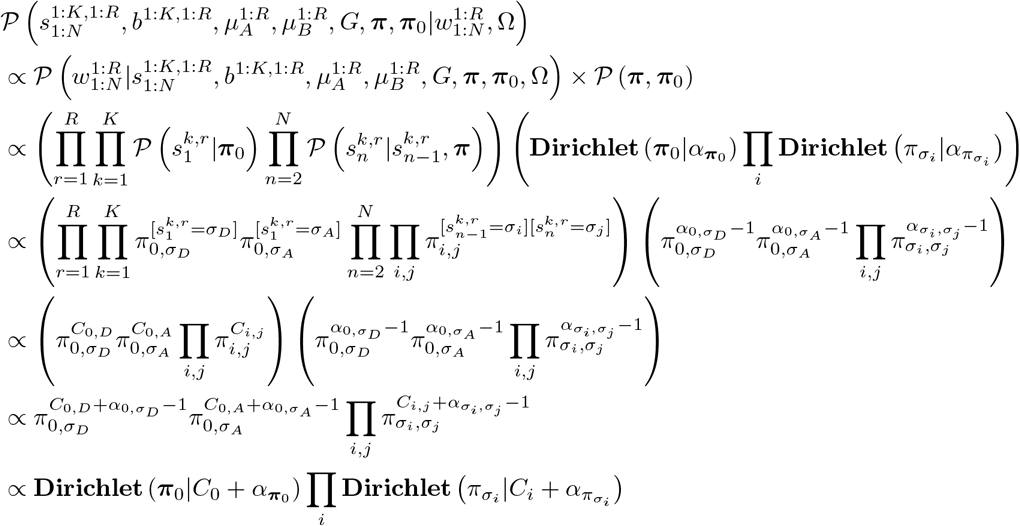

where *C*_0, *D*_ and *C*_0, *A*_ are the number of fluorophores that started in states *σ_D_, σ_A_*, and *C_i,j_* is the number of fluorophores that transitioned from *σ_i_* to *σ_j_*, *C*_0_ and *C_i_* are the arrays containing the counts *C_i,j_*, and the products indexed by *i* and *j* go over *σ_D_, σ_A_*, and *σ_B_*. So the Dirichlet prior on the initial state probability and transition probabilities allow us to get back independent Dirichlet distributions in the posterior.

Notice that all elements of ***π***_0_ and ***π*** that are equal to zero also have zero counts in the trace. By declaring 0^0^ = 1, these components do not affect the probability.

### 5.4 Hyperparameter on loads

Each load is sampled from a Bernoulli distribution with a parameter *q^k,r^*. This parameter *q^k,r^* is sampled from a beta distribution with hyperparameters *γ* and *K*, where *γ* is a hyperhyperparameter on all *q^k,r^*’s and *K* is the total number of load on and load off fluorophores in ROI *r*.

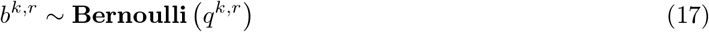

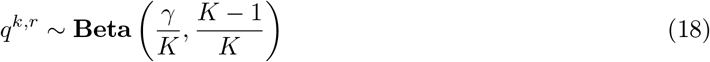

This is the Beta-Bernoulli process (BBP) [42, 43]. The distribution on *q^k,r^* is constructed in such a way that as *K* goes to infinity, *q^k,r^* goes to zero consistent with our assumption that *K* ≫ *K_r_*. Because *q^k,r^* is a hyperparameter, it is computationally advantageous for us to marginalize it out. We marginalize over the *q^k,r^*’s to get a Bernoulli distribution for *b^k,r^* in terms of *γ* and *K*.

*Proof (with indices dropped for clarity):*

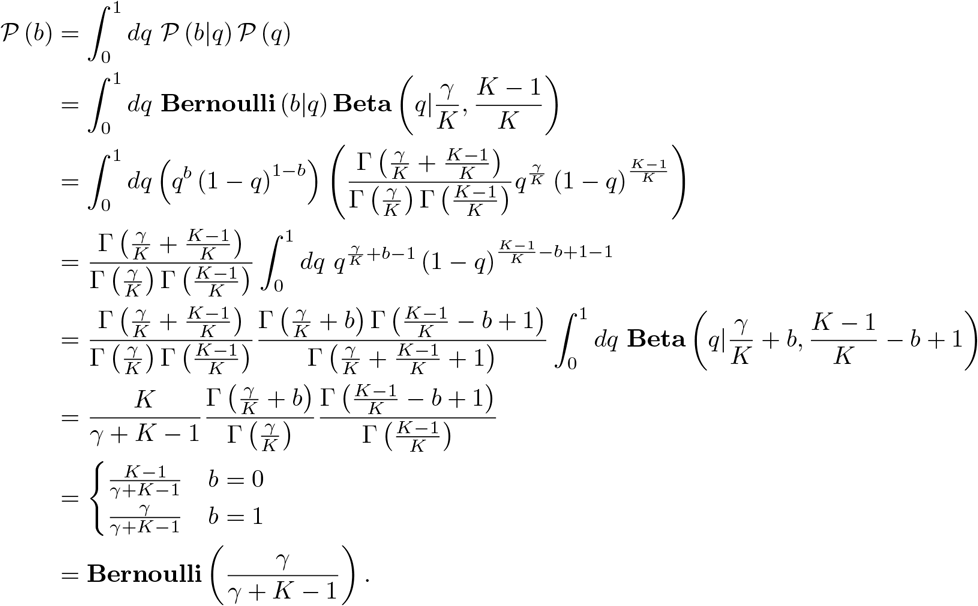

Following our assumption, *K* ≫ *K_r_, γ*/(*γ* + *K* – 1) must be a small number (we use *γ*/(*γ* + *K* – 1) = .1 with *K* = 100). Beyond this restriction, we can set *γ* by hand knowing that its effect is minimal for sufficient data. Altogether we have

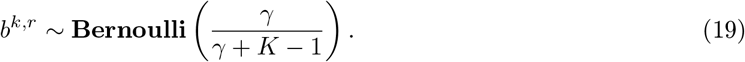

### 5.5 Multiple bright states

Here instead of having one active state, *σ_A_*, we will have *L* active states, *σ*_*A*1_,…, *σ_AL_* each with brightness *μ*_*A*1_,… *μ_AL_*. We note that *L* must be selected *a priori*. The transition matrix is now an *L* + 2 by *L* + 2 matrix

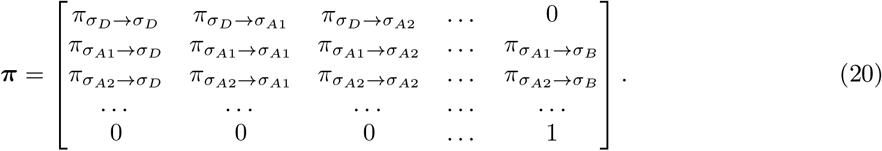

We note that as we add these bright states, we must add *L* different random variables (*μ*_*A*1_,… *μ_AL_*).

### 5.6 Model summary and posterior

The model is summarized as follows.

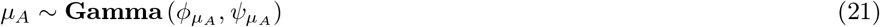

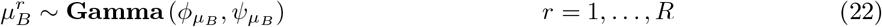

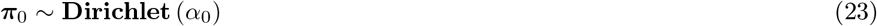

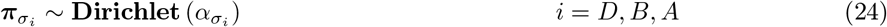

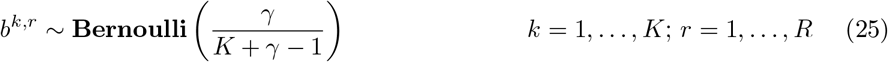

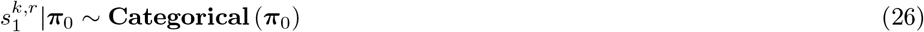

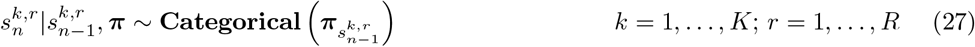

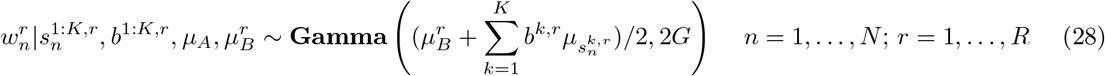

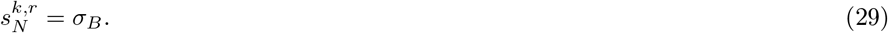

The inverse model scheme (section 2.2) allow us to develop the joint posterior of our model

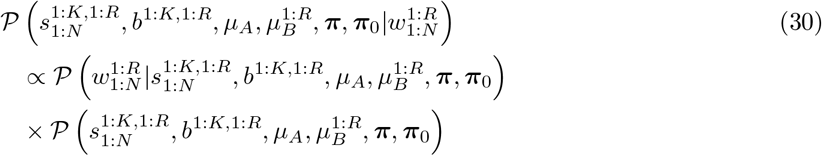

where the likelihood (middle term of Eq. (30)) is the product of Eq. (31) for all time levels and ROIs, which looks like

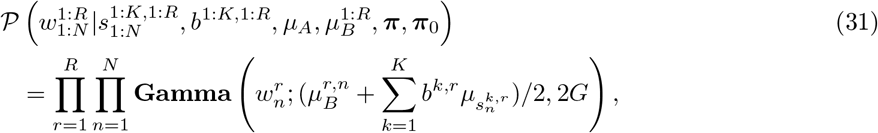

and the prior (final term) of can be further decomposed into

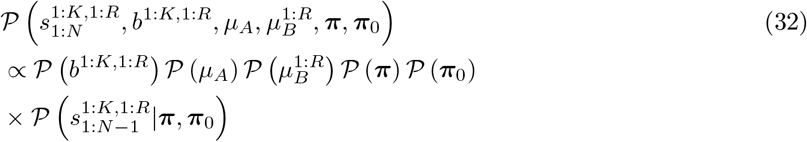

which has two main parts, 1) the middle terms of Eq. (32) are the product of Eqs. (11)–(6) over all indexes 2) the last term of Eq. (32) is the product of *π*_0,*D*_ to the power of the number of fluorophores that start dark (*C*_0*D*_ is the number of fluorophores that start dark) times *π*_0,*A*_ to the power of the number of fluorophores that start bright (*C*_0*A*_ is the number of fluorophores that start bright) times *π_ij_* to the power of the number of transitions that occurred from *σ_i_* to *σ_j_* (*C_ij_* is the number of transitions from *σ_i_* to *σ_j_*)

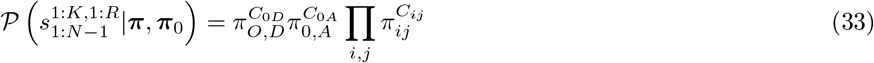

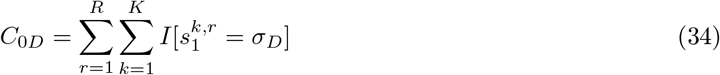

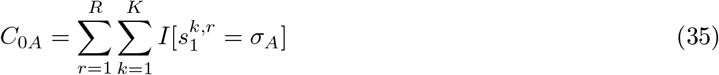

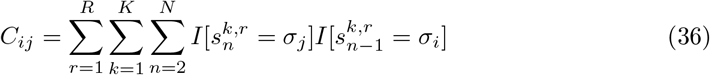

where *I*[*x* = *y*] is the indicator function that is equal to 1 if *x* = *y* and 0 else. We note that in this scheme (Eqs. (30)–(33)), we are learning the number of fluorophores in many ROI simultaneously. This is important because by analyzing many ROI together, we effectively have more information to train on and therefore obtain more accurate results.

### 5.7 Sampling

We outline the Gibbs sampling scheme below where (*i*) indexes the iteration in the Gibbs sampler.

- Step 1: Choose initial 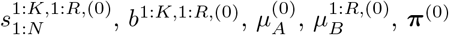, and 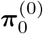.
- Step 2: For many iterations, *i*:

– A) Sample new states and loads from

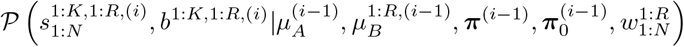

which is the product of Eq. (31) and Eq. (33).
– B) Sample new transition probabilities and initial state probabilities from

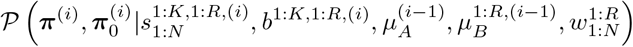

which is the product of Eqs. (11), (12), and (33). Here the choice of conjugate priors allow us to sample ***π***_0_ and ***π*** directly.
– C) Sample new camera parameters from

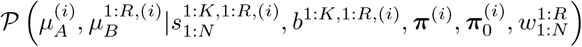

which is the product of Eq. (31) and the priors (Eqs. (13)–(14)).

For Step 1, the initial values are chosen to be the means of the prior. These means can in principle be anything, although, better guesses lead to faster convergence. We did notice the sampler is sensitive to the initial value for the fluorophore brightness. In other words we could sample, for example, half as many fluorophores with twice the brightness or three times the number of fluorophore with a third the brightness. To avoid this problem we set a sharp prior for the value of *μ_A_* on the brightness of a single fluorophore which is clear at the end of the trace. This makes sure that our model posterior is sharply peaked at reasonable numbers of fluorophores. We calibrate gain, *G*, using methods described in Hirsch et al [44]. We estimate the background brightness for each ROI by looking at the end of the brightness time trace, when all fluorophores are presumed to be photobleached. For 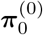 we take the maximally naive approach and choose the starting probability for bright and dark to be 50/50. We can choose **π**^(0)^ using this approach, but we find that choosing self transitions to be larger speeds up the convergence significantly (by about 100-500 iterations) For 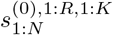 we guess that there are no fluorophores in any ROI, i.e that for all loads, *b^k,r^* = 0. In our analysis, we chose the hyperparameters to coincide with the initial guess. So *α*_0_ = [.5, .5,0], *α_σ_D__* = [.9, .1,0], *α_σ_A__* = [.1, .8, .1], *α_σ_B__* = [0, 0,1]. For the mean background brightness and mean fluorophore brightness, we chose the initial guess to be the mean of the prior by scaling the scale parameter by the shape parameter,

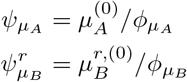

In Step 2 A, for each ROI, we sample loads and states together by using the forward filter backwards sample algorithm [48]. To speed up computation we do this using a collapsed state space where we treat photobleached fluorophores and fluorophores with load off to be the same. That is, under our model each fluorophore is described by two different random variables, its load and its state. However, we instead describe it with one random variable by lumping together fluorophores with off loads and photobleached fluorophores with on loads. Consider a new state space defined by

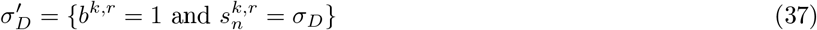

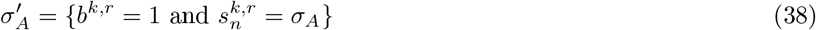

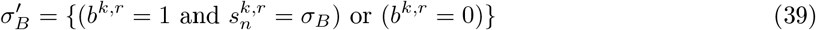

In this “collapsed state space”, we ignore the loads entirely, but allow for fluorophores to start from the photobleached state with probability (*K* – 1)/(*γ* + *K* – 1) (the probability that a fluorophore has load off). This in turn causes ***π***_0_ to scale down by *γ*/(*γ* + *K* – 1) as well,

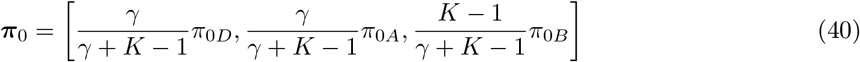

but notice that ***π*** is not affected (as off load fluorophores cannot become on and vice versa) nor are any of the other random variables. Thus we can simplify our calculations by simply allowing load off fluorophores to be considered photobleached so long as we keep track of ***π***_0_. Additionally, rather than sample each 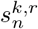 individually, we sample 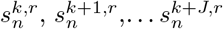 together (for *J* = 4 in our case). We additionally shuffle the fluorophore indices *k* so that at each sampler iteration, different fluorophores are sampled together. Sampling fluorophore states together in this way helps the sampler mixing. The joint microstate, 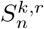, specifies the states of the fluorophores when we consider them together. For example, one realization of a joint microstate might look like

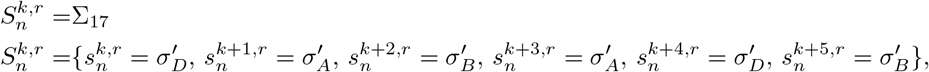

or using a notation Σ_1_, Σ_2_,… to represent the possible joint microstates that 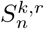 can take,

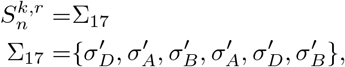

where the index, 17, on Σ_17_ is used simply to illustrate an example. The number of values 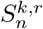 can take (Σ_1_, Σ_2_,…) is the Kronecker product of the individual state spaces 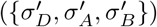. For Step 2 C, we sample *μ_A_* and 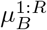 each separately using a Hamiltonian Monte Carlo step [66].

As the sampler runs, higher probability regions are sampled more often and lower probability regions are sampled less often [51, 52, 40, 53]. After many iterations we can histogram the results to approximate the shape of our posterior. By looking at the histogram and not just the MAP or mean, we are able to get credible intervals for our estimates.

We note that the *R* ROIs in the base set are analyzed simultaneously. To be clear, we mean that each ROI has its own fluorophore states, loads, and mean background brightness, but the transition probabilities, initial state probability, and mean fluorophore brightness are global. This is advantageous because it allows the inference of each ROI to benefit from the others by providing more data on which to train the transition probabilities.

### 5.8 Hamiltonian Monte Carlo

We sampled our state brightness and background brightness using Hamiltonian Monte Carlo [66]. Briefly, Hamiltonian Monte Carlo proposes samples using a numerical integrator with strong parallels to Hamiltonian dynamics. Here, our random variables are sampled along with normally distributed auxiliary variables, referred to as momenta, *p_A_* and *p_B_*

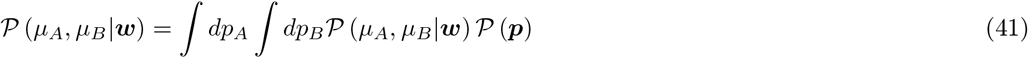

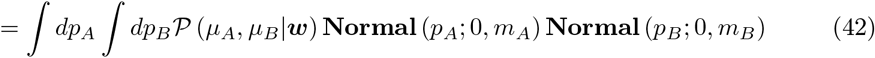

where *m_A_* and *m_B_*, called the masses, are the variances of the priors on the momenta (for simplicity in the derivation we only look at a single bright state and single background brightness). The negative log of the conditional posterior, including the momenta, is called the Hamiltonian.

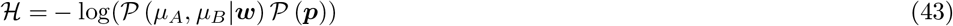

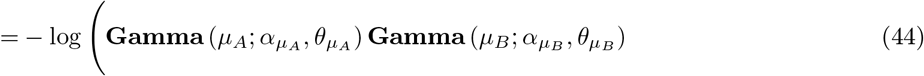

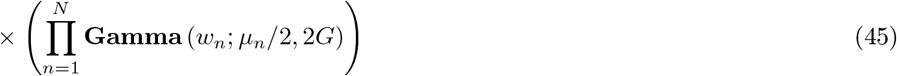

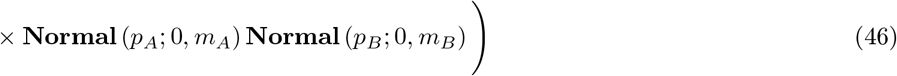

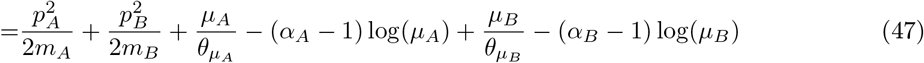

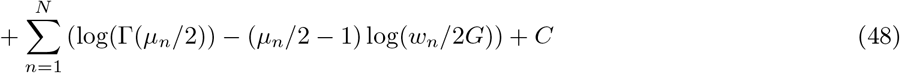

where *μ_n_* = *μ_B_* + *X_n_μ_A_* with *X_n_* is the number of fluorophores in the bright state at time level *n*, and where *C* includes all terms not dependent on *p_A_, p_B_, μ_A_*, or *mu_B_*. Note that the momenta in the Hamiltonian appear exactly like kinetic energy terms (motivating the name Hamiltonian Monte Carlo). We may use Hamilton’s equations of motion to modify *p_A_, p_B_, μ_A_*, and *μ_B_* in such a way that the total “energy” of the system is conserved

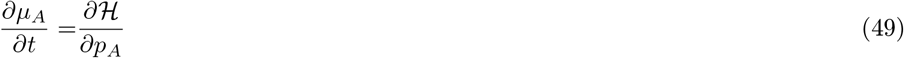

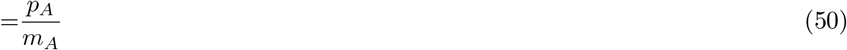

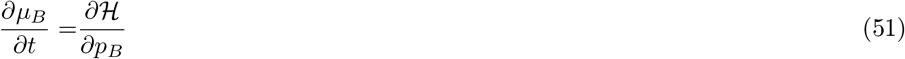

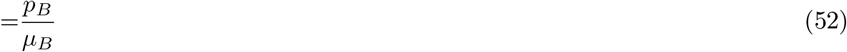

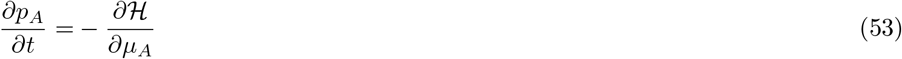

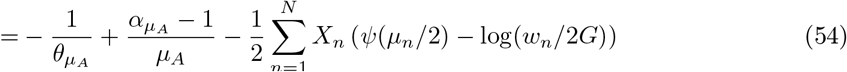

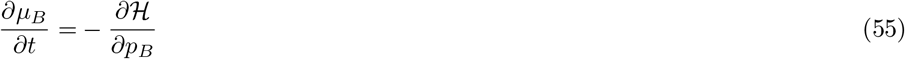

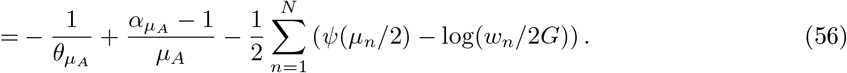

Conservation of energy in this case corresponds exactly to conservation of probability in the posterior. In principle, this means that starting with sampled values for *p_A_* and *p_B_*, we can use a Hamiltonian mechanics integrator to find proposal values of *μ_A_* and *μ_B_* with equal probability as those from the previous Gibbs sampler iteration.

### 5.9 Ruler method

The ruler method estimates the number of fluorophores by dividing the initial brightness by the estimated fluorophore brightness. For our implementation of the ruler method we average the last 10% of the brightness trace to get an estimate for the background brightness and we average the first 10 data points to get an estimate for the initial brightness. To estimate the brightness of a single fluorophore we average the last ten frames of the final photobleaching step less the background. The final photobleaching step is found by looking for the last time level where the brightness is above a set threshold. For our purposes, this threshold is always chosen to be the same brightness guess that we use to inform our prior on brightness, except for the high noise experiment (figure 4 bottom row) in which the high noise causes the background brightness to be frequently measured higher than the fluorophore brightness step. For the high noise case, we set the threshold to be 3 times the fluorophore brightness guess, which is high enough that the background is excluded, but low enough that it is sampled in the final photobleaching step (see figure 5.13). We note that setting the brightness of a fluorophore by hand can improve the ruler method’s accuracy. However, as our method, the change point method, and the two state method simultaneously learn fluorophore brightness with the number of fluorophores, we require that the ruler method also learn the fluorophore brightness in establish a fair comparison.

### 5.10 Robustness analysis on simulated data

In this section we evaluate the robustness of our method. In the main (section 3.2), we demonstrated the method using a “base set of parameters”. For the base set we have 50 ROIs containing 14 fluorophores on average. The traces are 1000 seconds long with brightness 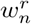 collected every 50 *ms*, so 20000 total frames. The exact number of fluorophores in each ROI is sampled from a binomial distribution to mimic 20 binding sites with 70% labeling efficiency. The gain used for the simulation was 50. The dimensionless background brightness parameters are 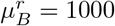. The fluorophores were simulated with two bright states with brightness given by *μ*_*A*1_ = 450 and *μ*_*A*2_ = 350 (plus one dark state and a photobleached state with brightness given by *μ_D_* = *μ_B_* = 0). For the number of loads, *K*, we chose 39. These parameters were chosen to mimic the parameters learned from real data, for example, the brightness states (*μ*_*A*1_ and *μ*_*A*2_) were chosen so that the height of a photobleaching step in the simulated data is the same height as a photobleaching step in the real data. We will refer to these parameters as the “base set.”

#### 5.10.1 Varying the number of loads

We must make sure that our choice of nonparametric limit in our beta-Bernoulli prior, Eq. (6), does not heavily the number of loads we choose. That is, we want to make sure that if 14 fluorophores are present in an ROI, then our choice of *K* = 30 or *K* = 300 from Eq. (6) is inconsequential. To address this we ran inference on the same simulated data, but with a different number of loads. The results are shown in figure 6.

**Figure 6:**
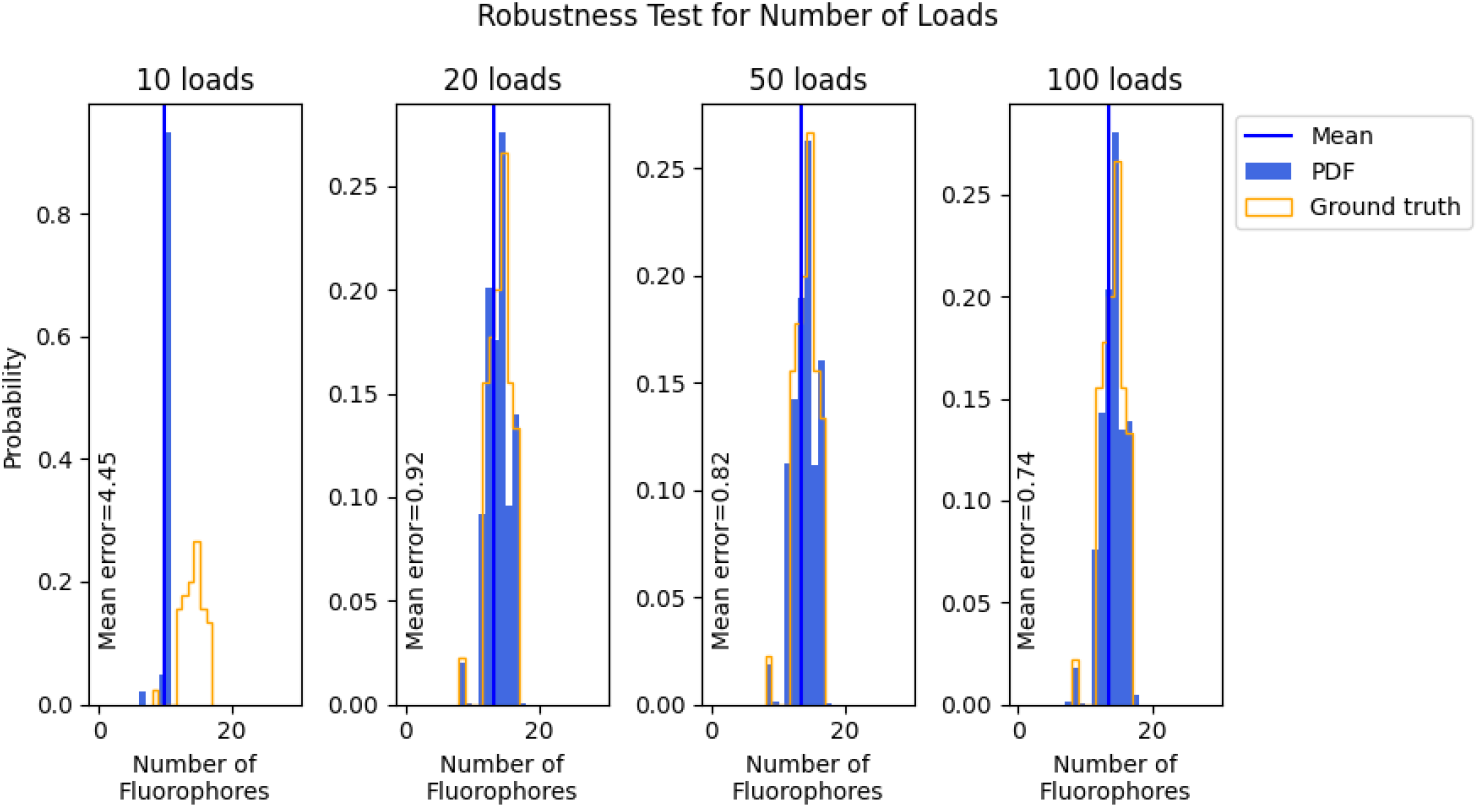
Inference on fluorophore number from data simulated with varying number of loads. Each panel shows inference on simulated data using a different number of loads. The mean error is annotated on the bottom left of each panel.

The top row of figure 6 shows that if we do not provide a sufficient number of loads, then we cannot count the correct number of fluorophores. This is obvious because if the number of fluorophores is determined by how many loads are “on”, then when there are not enough loads, we cannot turn on enough loads to account for all the fluorophores. For example, if we provide 9 loads, but there are 14 fluorophores, then even if all 9 loads are “on” we underestimate by 5 fluorophores. Aside from this constraint, we found that once a sufficient number of loads are chosen then the results are not dependent on the number of loads. As seen in the bottom three rows of figure 6, increasing the number of loads has negligible impact on the performance of the sampler.

Therefore, for the rest of this paper we will take for granted that we are free to set the number of loads, *K*, as we please provided that we set *K* larger than the maximum expected number of fluorophores. A note however is in order. If we choose too many loads, then the computation becomes expensive and inefficient. As a nice balance between sufficiency and efficiency, we will choose to make the number of loads equal to two times the expected number of fluorophores. In the case that the ground truth number of fluorophores is unknown (like for real data) the best way forward would be to guess a number of loads and if it maxes out (i.e., the number of sampled fluorophores in an ROI is equal to the number of loads) then to double the number of loads and try again.

#### 5.10.2 Varying the number of fluorophores

Now that we have shown that we can accurately count the number of fluorophores in an ROI, the next clear step forward is to see how high we can count. For this, we simulated data using the same base case parameters, but changed the way in which we sample the ground truth number of fluorophores. Here we simulate ten different experiments with 20, 40,…, 200 fluorophore binding sites and 50% labeling efficiency so that the expected number of fluorophores for each ROI is 10, 20,…, 100 respectively.

As seen in figure 7, on simulated data for the parameters chosen, the learned distribution of fluorophores overlaps with the ground truth for all simulations. There is a slight underestimation, in which the sampler misses about 1 fluorophore out of every ten, that becomes more noticeable as the number of simulated fluorophores increases. The source of this underestimation is the prior on the number of loads(equation 6), which favors samples from the posterior that use fewer fluorophores to explain the data. At the beginning of the brightness trace, where the width of the brightness levels is much wider than the separation between brightness levels (figure 8), the sampler may “choose” to interpret steps as noise fluctuations, thus missing fluorophores. This underestimation percentage is consistent across number of fluorophores used for the simulations (we have similar error percentage when there are 20 ground truth fluorophores as when there are 100 fluorophores) and thus does not negate that we can count high number of fluorophores.

**Figure 7:**
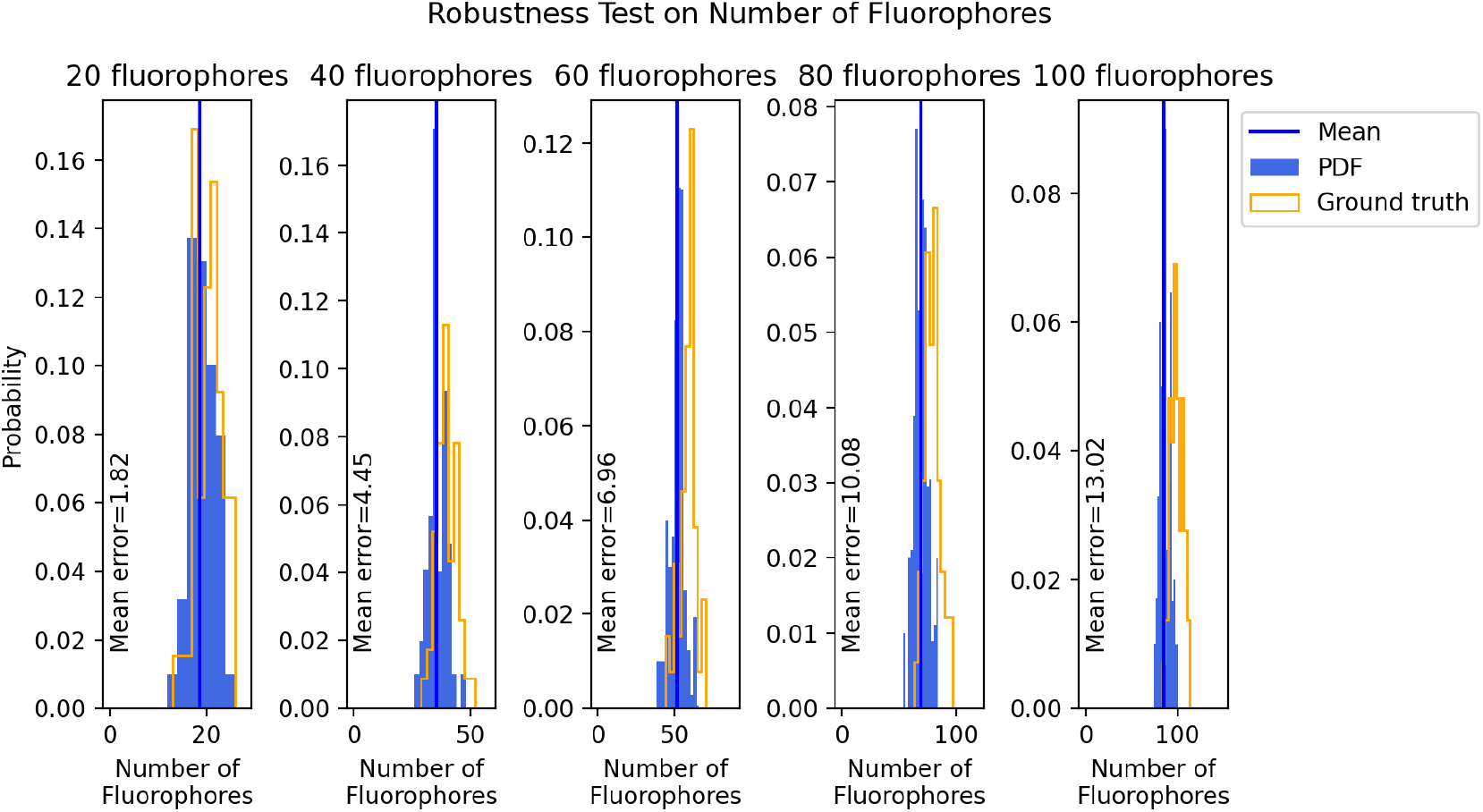
Inference on fluorophore number from data simulated with varying fluorophore numbers. Each panel shows inference on data simulated with a different number of fluorophores per ROI.

**Figure 8:**
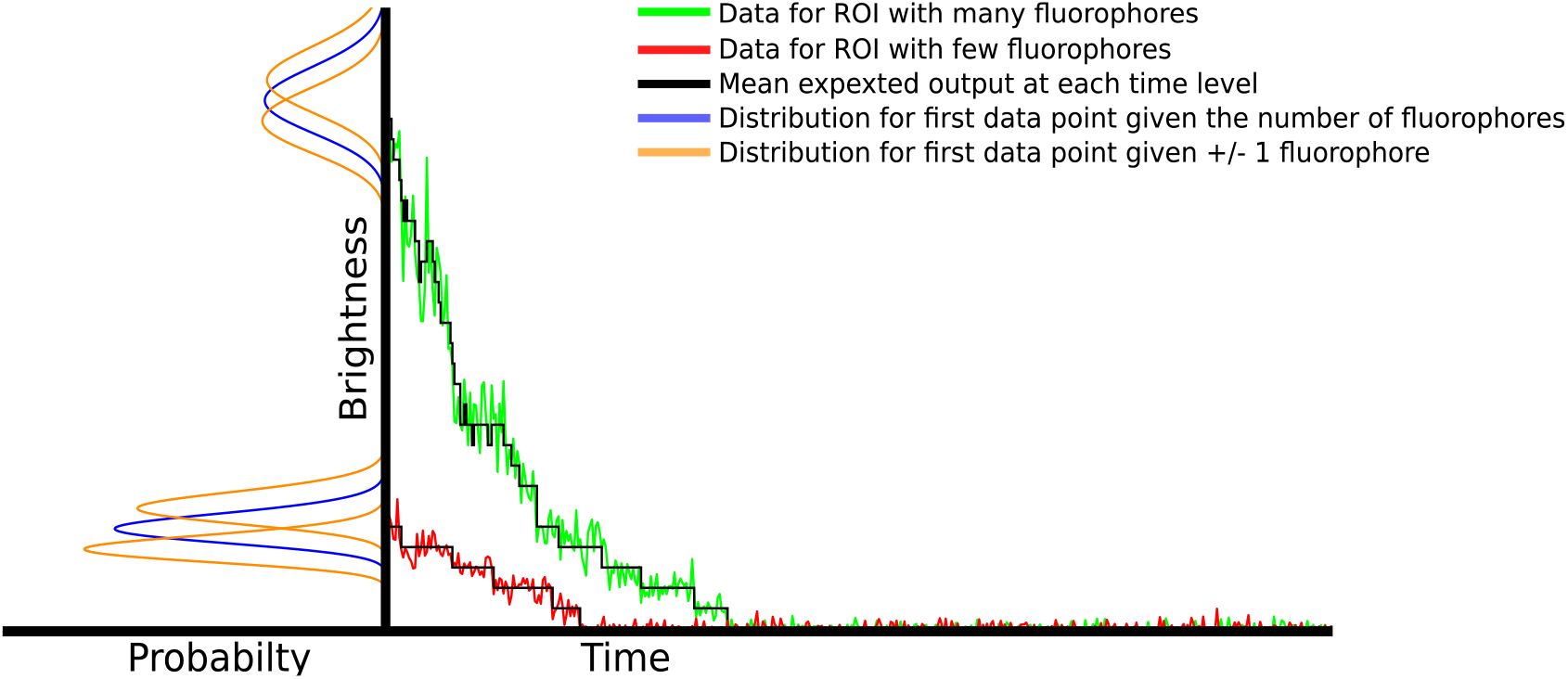
As the number of active fluorophores increases, photon shot noise, amplified by camera noise, becomes an essential modeling component. On the right we see two brightness time traces for an ROI. The ROI giving rise to the red trace contains 5 fluorophores and the ROI giving rise to the green trace contains 26 fluorophores. The black line through each trace represents the expected mean brightness at each time level based on the background brightness, fluorophore brightness, and number of active fluorophores at the time level. On the left we see the probability distribution for the initial brightness of each ROI (blue) as well as the probability distribution for the initial brightness if the ROI contained one more or fewer fluorophores (orange). We notice that: 1) for the red trace, the measured brightness matches closely with the mean, whereas the green trace fluctuates wildly; 2) the probability for the initial brightness is sharply peaked for the red trace which allows us to easily tell how many fluorophores are initially active. By contrast, the probability for the initial brightness is much more spread out for the green trace and overlaps greatly with the distributions for one more and one fewer fluorophores.

#### 5.10.3 Varying the number of states

We note that while our model is nonparametric in the number of fluorophores in an ROI (i.e., that the number of fluorophores in an ROI is not a fixed quantity but inferred), our model is parametric in the number of fluorophore states. That is, we must pre-specify the number of fluorophore states. It is difficult to be both nonparametric in the number of fluorophores and the number of states (as then, trivially, each transition could be considered to visit a new state of the fluorophore). In the previous sections, we have taken for granted that we knew the correct number of fluorophore states. We modeled that the fluorophores have two separate bright states, a dark state, and a photobleached state. Here we explore the photophysical state space further using different numbers of bright states. For conciseness, in this section we will refer to these models using a naming convention that starts with ‘D’ followed by the number of dark states and then ‘B’ followed by the number of bright states: 1) D0B1, a model with no dark state and one bright state (i.e., the fluorophore is either bright or photobleached); 2) D1B1, a model with one dark state one bright state; and 3) D1B2, a model with one dark state and two bright states. In principle we could also explore more bright or dark states, but the combinatorics on the possible arrangements on fluorophore states adds computational burden. We note that the base case set of parameters uses a D1B2 model and the model with only two states, D0B1, is the same model as the one used in Garry et al. [25].

We simulated data using each of the different state models and then analyzed each simulated data set using each of the different state models. Results are shown in figure 9. Starting left to right, we first notice that the D0B1 model does well at counting fluorophores from data generated using the D0B1 model (figure 9 top left). On the other hand, the inference severely underestimates the number of fluorophores for the data generated using the D1B1 model (figure 9 middle left) and the D1B2 model (figure 9 bottom left). This is because, lacking a dark state, no possible fluorophore state trajectory under the D0B1 model can account for blinks where the brightness of the ROI goes down and then comes back up. Thus the only way for the method to account for blinks is to increase the variance (by increasing the background brightness, *μ_B_*) to a level where a sudden drop in brightness is explained purely by noise. This in turn makes photobleaching steps the same size as noise fluctuations (because the drop in brightness from a blink is the same as a drop in brightness from a photobleaching event), thus the method “misses” photobleaching steps and most brightness drops are explained purely by the exponentially decaying background.

**Figure 9:**
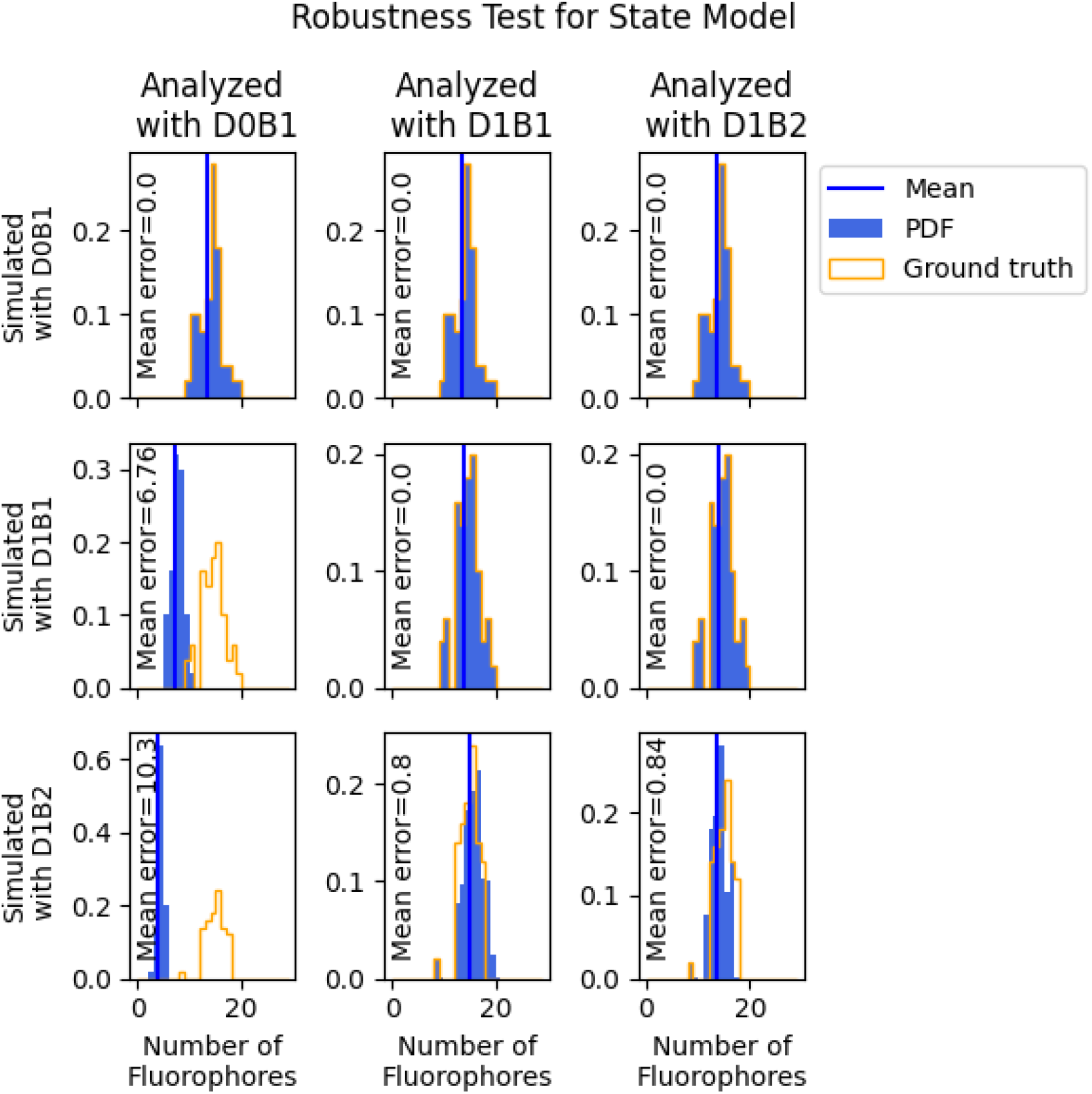
Inference on data simulated with different number of states. Here for each row we generate simulated data using a different state model for the forward model. Each column shows the inference on that data set using a different state model for the inverse model.

The remaining models, D1B1 and D1B2, provide good matches to the ground truth number of fluorophores for the data with no dark state (figure 9 top row). In this case the dark state is never visited despite being available. Similarly both models are able to exactly learn the ground truth number of fluorophores in the data sets with only one bright state. Again, in this case the two bright state inverse model will simply rarely visit one of its available bright states. For the data generated using two bright states, both the single bright state model, D1B1, and the two bright state model, D1B2, both infer a distribution of fluorophore numbers with similar accuracy. As the two bright states are very close to each other in brightness, the D1B1 model is able to find a brightness that averages the two together. For this reason, we say that our inference scheme is robust to bright state number. The advantage to using the two bright state model, D1B2, is that the inferred brightness traces will learn bright state transitions kinetics.

### 5.11 Additional comparison

Here we plot additional figures comparing our method to the change point detection algorithm laid out in Tsekouras et al. [15] and the ruler method[12, 13].

In figure 11 we plot the learned distribution for the number of fluorophores for each method as well as the ground truth distribution for the number of fluorophores. On the bottom left of each panel we show the mean error of each method, that is the average difference between the predicted number of fluorophores and the ground truth number of fluorophores for each ROI. Our method provides the least error for all data sets except the 80 fluorophore data set in which the ruler method has less error. This is likely a statistical artifact in which the noise fluctuations at the beginning of the trace and end of the last photobleaching step made estimating the fluorophore brightness and initial brightness easier. Note on the other hand that because such statistical deviations are rare, for every other data set our method is more accurate than the ruler method for this point statistic alone (determining fluorophore counts).

**Figure 10:**
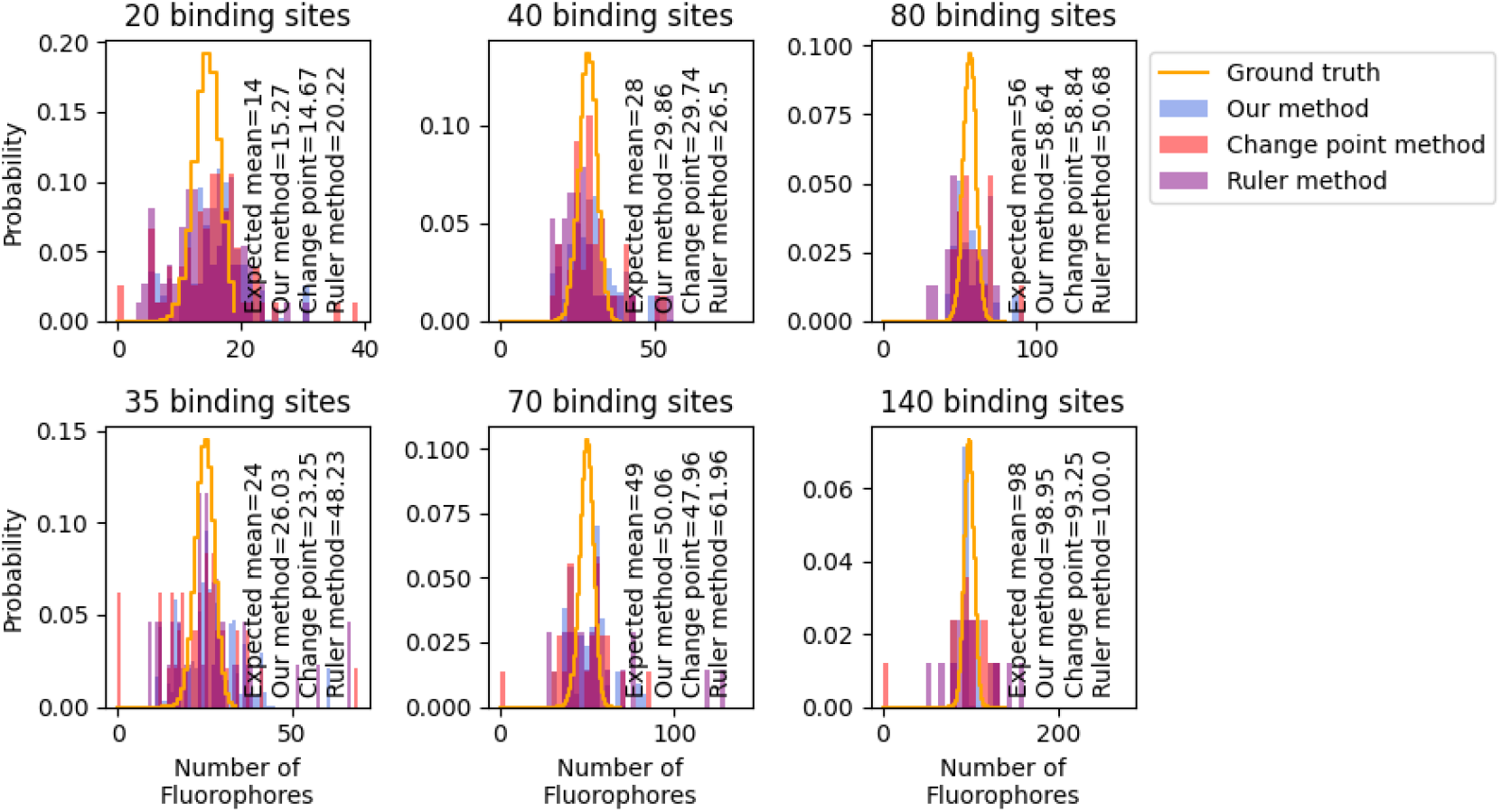
Comparison against other methods on real data. Here we compare our method against a change point method and the ruler methods on real data. Each panel shows the inference on a data set with a different number of fluorophores. We plot the inferred distribution for the number of fluorophores for each method in a different color. On the bottom right of each panel we annotate the mean expected number of fluorophores as well as the learned mean expected number of fluorophores.

**Figure 11:**
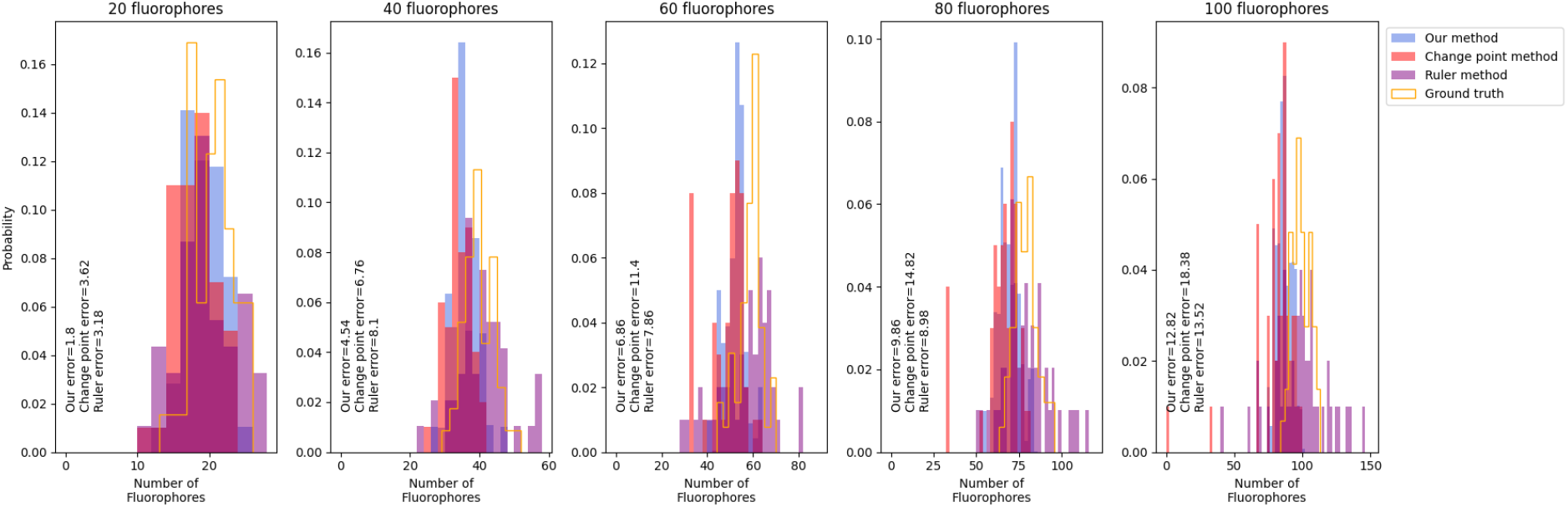
Additional comparison against other methods on simulated data. Here we compare our method against a change point method [15] and the ruler methods on real data. Each panel shows the inference on a data set simulated with a different number of fluorophores. We plot the inferred distribution for the number of fluorophores for each method in a different color. On the bottom left of each panel we annotate the mean error.

In figure 10 we plot the learned distribution for the number of fluorophores for each method as well as the ground truth expected distribution for the number of fluorophores. As there is no known ground truth, we cannot compare errors. Instead we compare the learned average number of fluorophores against the best guess for the expected average number of fluorophores given the factory estimated binding rate. We note that the ruler method occasionally predicts very high number of fluorophores (200+) when the fluorophore brightness is underestimated due to blinking or random spikes.

We can use our obtained mean number of fluorophores to validate the 70% binding efficiency provided to us by the manufacturer. We do this by dividing the learned mean number of fluorophores by the number of binding sites. Doing this we estimate 76% binding efficiency from our 20 binding site data and we estimate 74% binding efficiency from our 35 binding site data. Both of these estimates are within 10% of the provided estimate.

### 5.12 Full posteriors

Here we plot full posterior samples for each unknown quantity from the analysis on each data set. We do this in two figures for each experiment. For the first part we plot the Gibbs sample trace showing the values of each of our parameters as a function of iteration. The top panel of each figure shows the log probability at each iteration. The second panel shows the brightness of the bright states at each iteration. The third panel shows the lifetime of the bright states at each iteration. The final panel shows the learned average number of fluorophores at each iteration, that is, we learn the number of fluorophores in each ROI and then average them. For simulated data sets we plot the ground truth values as a horizontal line. As there are two bright states, the middle two panels will have two Gibbs sample traces (there are also two ground truth values, but they are very close together). For the second figure we show a scatter plot of the covariance of the posterior. For this plot, each row and column represent a different variable (we simplify by only showing the brightness of the brightest state, the lifetime of the brightest state, and the mean number of fluorophores per ROI). Each panel *i, j* shows there covariance between variable *i* and variable *j* (where, again, *i* and *j* will be the brightness of the brightest state, the lifetime of the brightest state, and the mean number of fluorophores per ROI) and each panel along the diagonal shows the histogrammed posterior of variable *i*.

The input parameters used for analysis are the same for each data set and correspond to the default inputs on the provided code. We reproduce those default values below. Let *u_r_* be the average of the last 100 data points of ROI *r* which is used for estimating the background brightness.

|  |  |  |
| --- | --- | --- |
| $f$ | fluorophore brightness guess | 10000 |
| $G$ | gain* | 22 |
| $\gamma$ | hyperparameter on loads | .1 |
| $\phi_{\mu_A}$ | scale parameter for fluorophore brightness | 100 |
| $\psi_{\mu_A}$ | shape parameter for fluorophore brightness | $f/(G \times \phi_{\mu_A})$ |
| $\phi_{\mu_B}^r$ | scale parameter for background brightness | 10 |
| $\psi_{\mu_B}^r$ | shape parameter for background brightness | $u_r/\phi_{\mu_B}^r$ |
| $\pi$ | concentration parameter for transitions | $\begin{bmatrix} 100 & 1 & 1 & 0 \\ 1 & 100 & 1 & 1 \\ 1 & 1 & 100 & 1 \\ 0 & 0 & 0 & 1 \end{bmatrix}$ |
| $\pi_0$ | concentration parameter for initial state | $\begin{bmatrix} 1 & 1 & 1 & 0 \end{bmatrix}$ |

\*Gain is $G = 2200$ for the experiment in figure 4 bottom row.

For all experiments we perform one MCMC chain with a set random seed. We run our Gibbs sampler for 20,000 iterations or 1 week, whichever comes first, then eliminate the burn in phase by selecting only the samples where the variables and the log posterior have stabilized. Our heuristic metric for stablization is when the slope of all samples with respect to Gibbs sample iteration is less than one percent of the mean.

Note that the state lifetimes are calculated from the transition learned transition matrix as a post processing step via *τ_k_* = dt/(1 – *π_k,k_*) where *τ_k_* is the lifetime of state *k* and *π_k,k_* is the self transition probability of state *k*. Under this transformation, small displacements in *π_k,k_* can cause large displacements in *τ_k_*. For this reason, the learned bright state lifetimes can be off by as much as a factor of 2 even if the learned transition rate is only off by 10% or less.

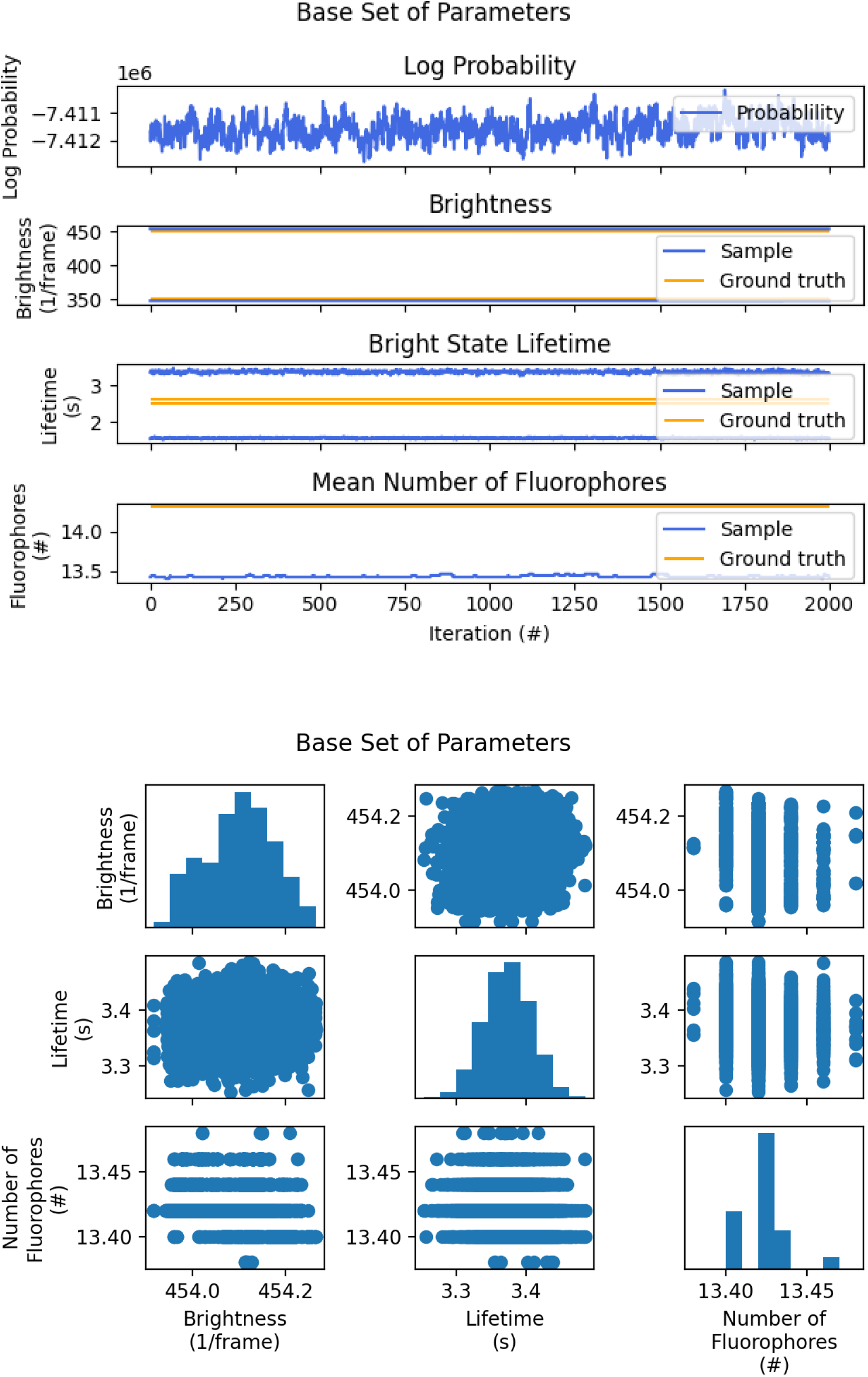

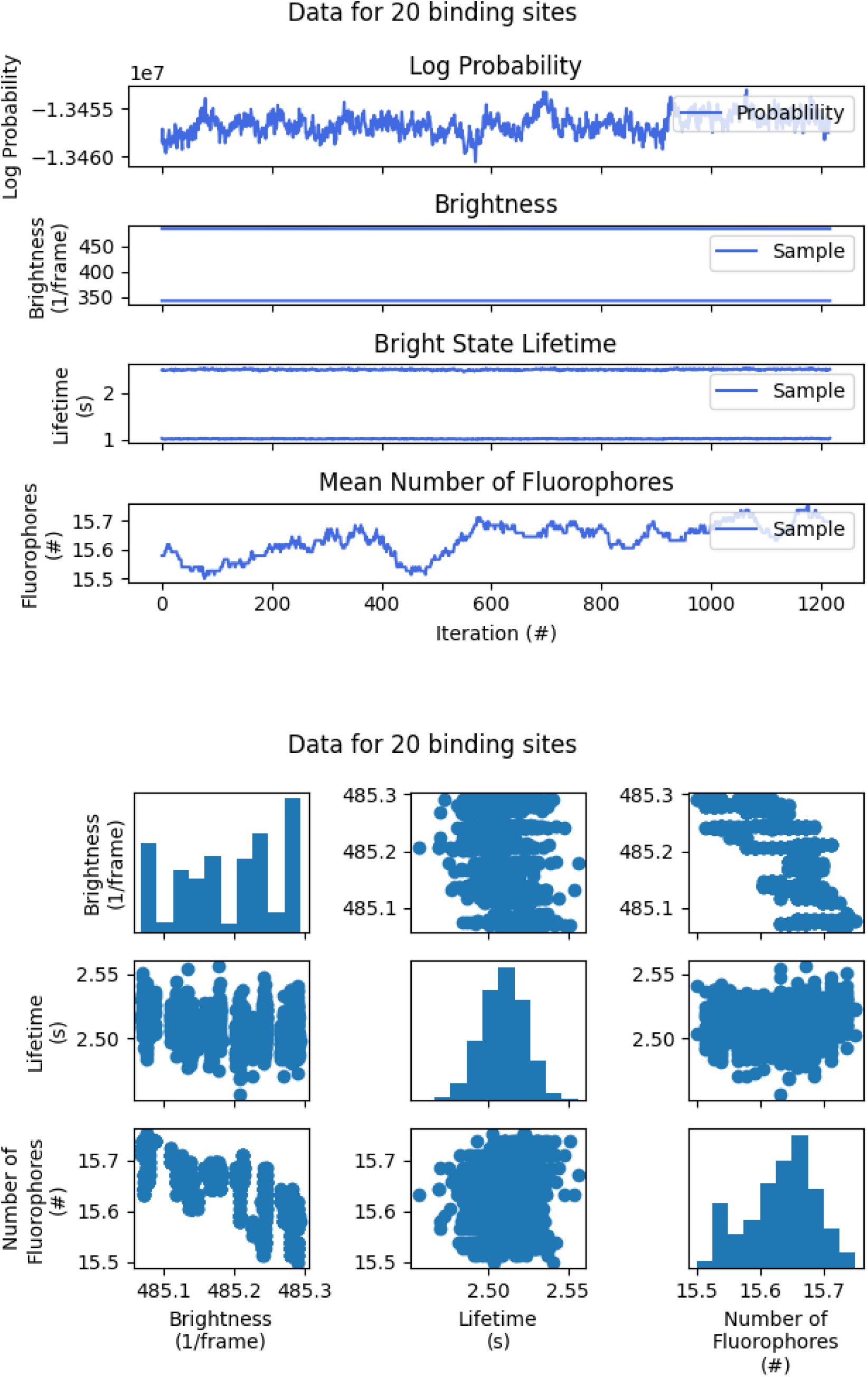

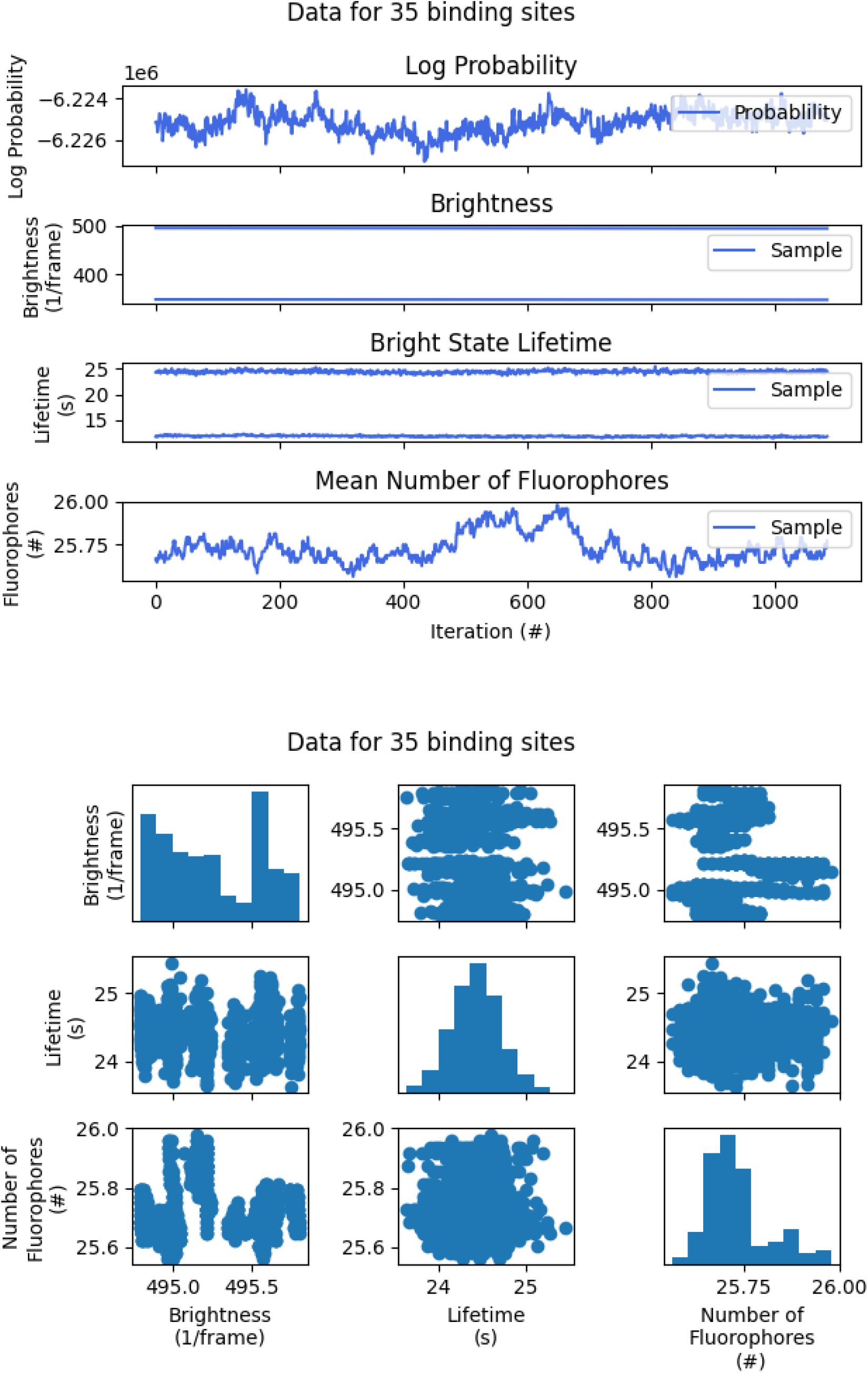

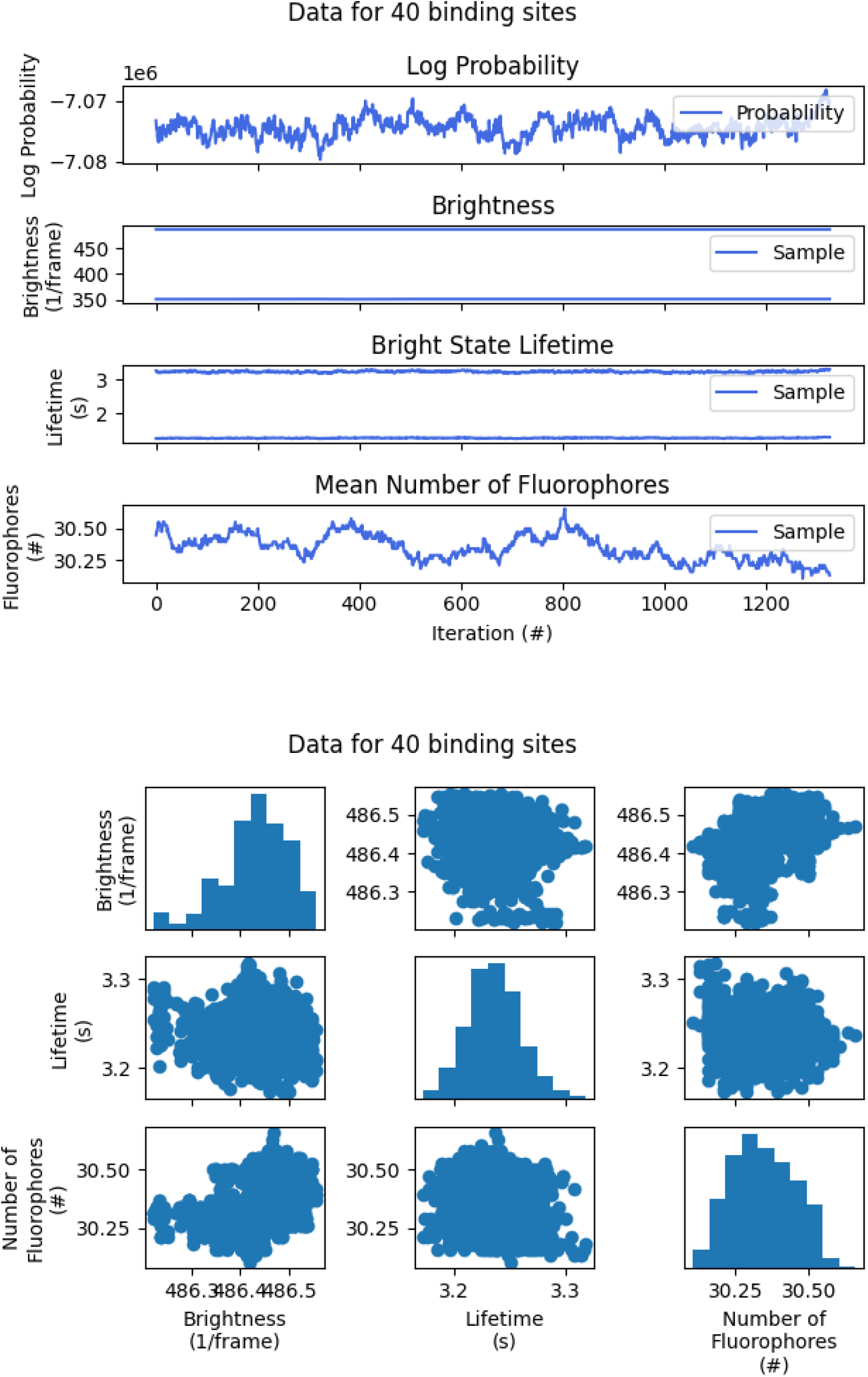

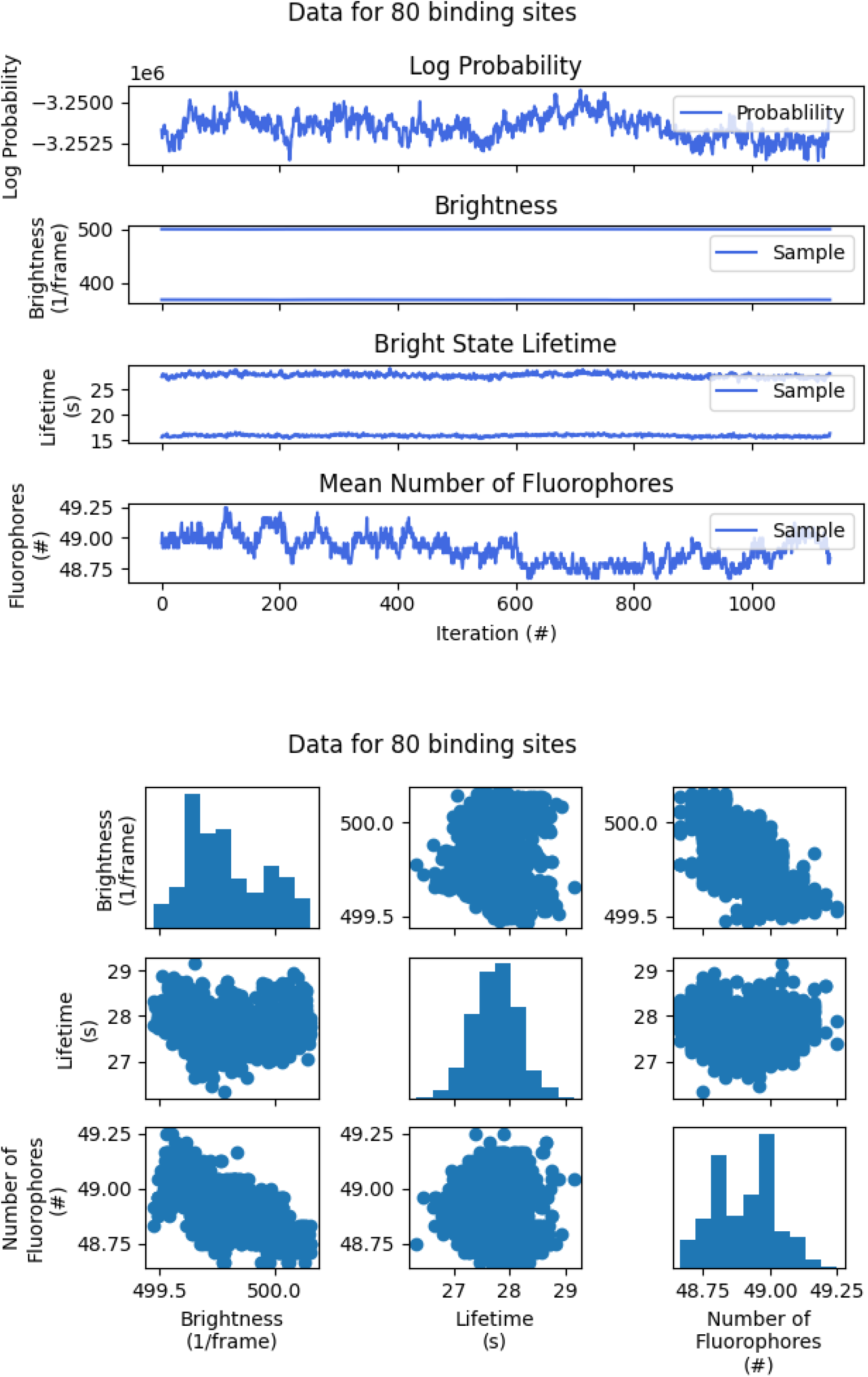

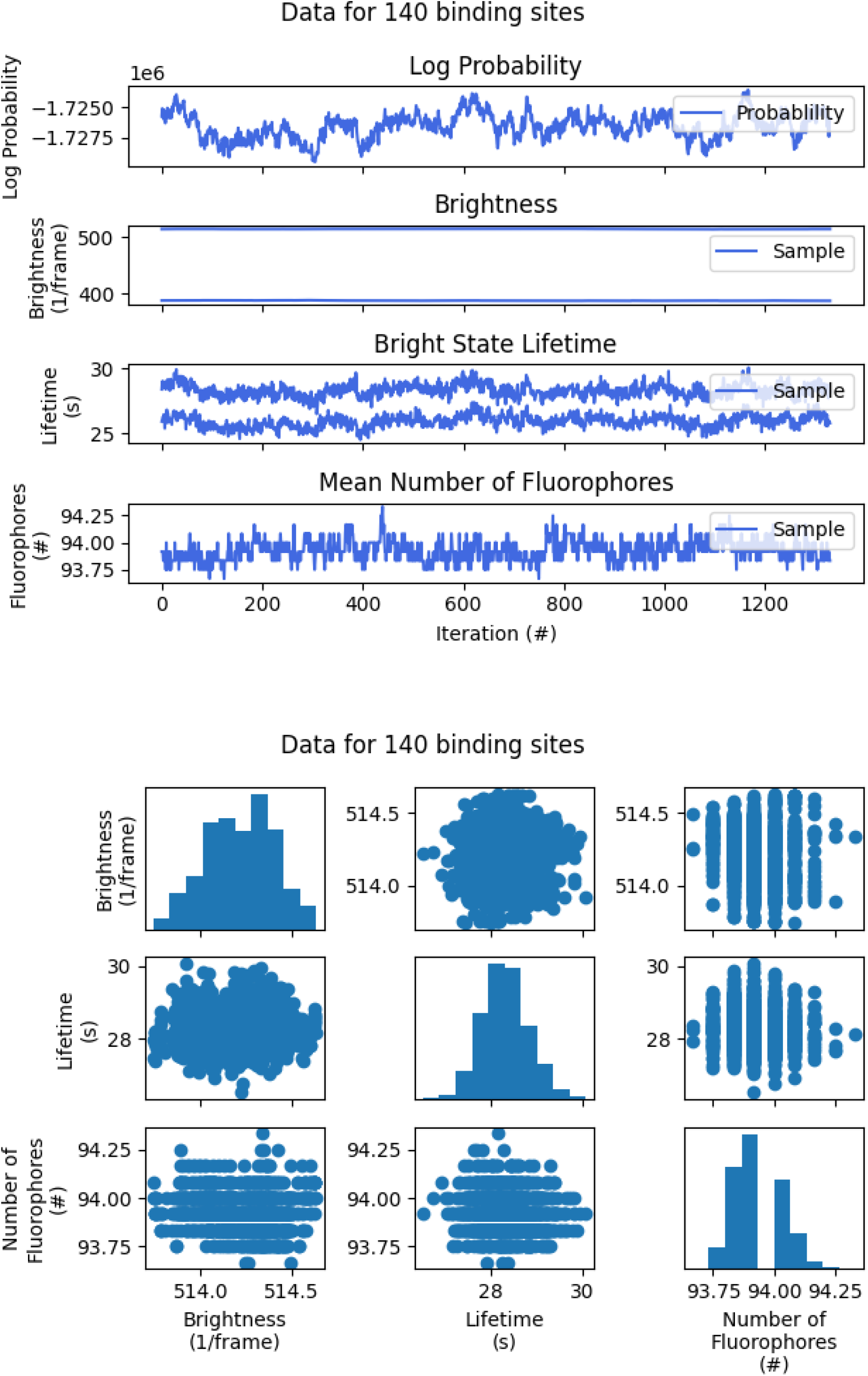

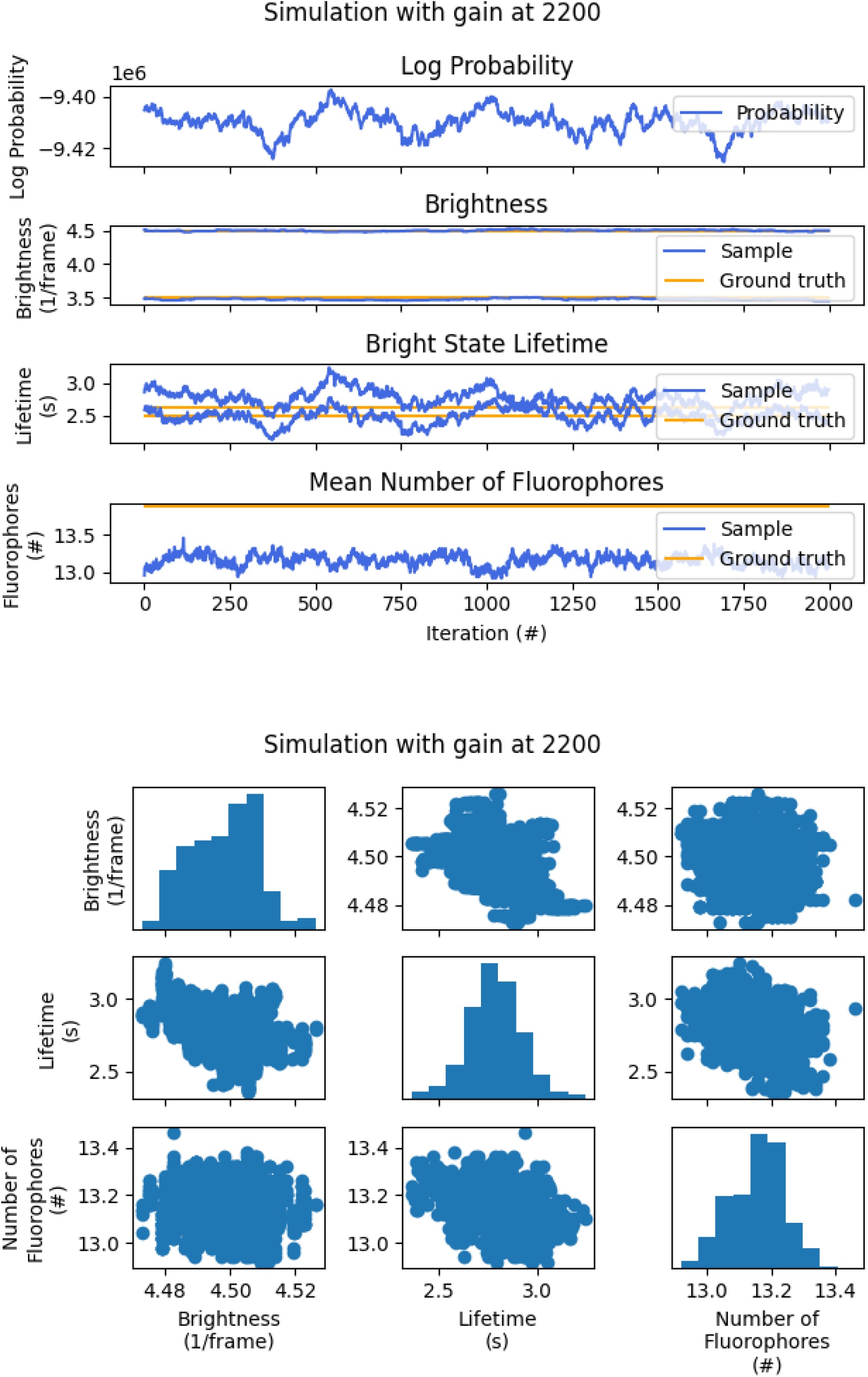

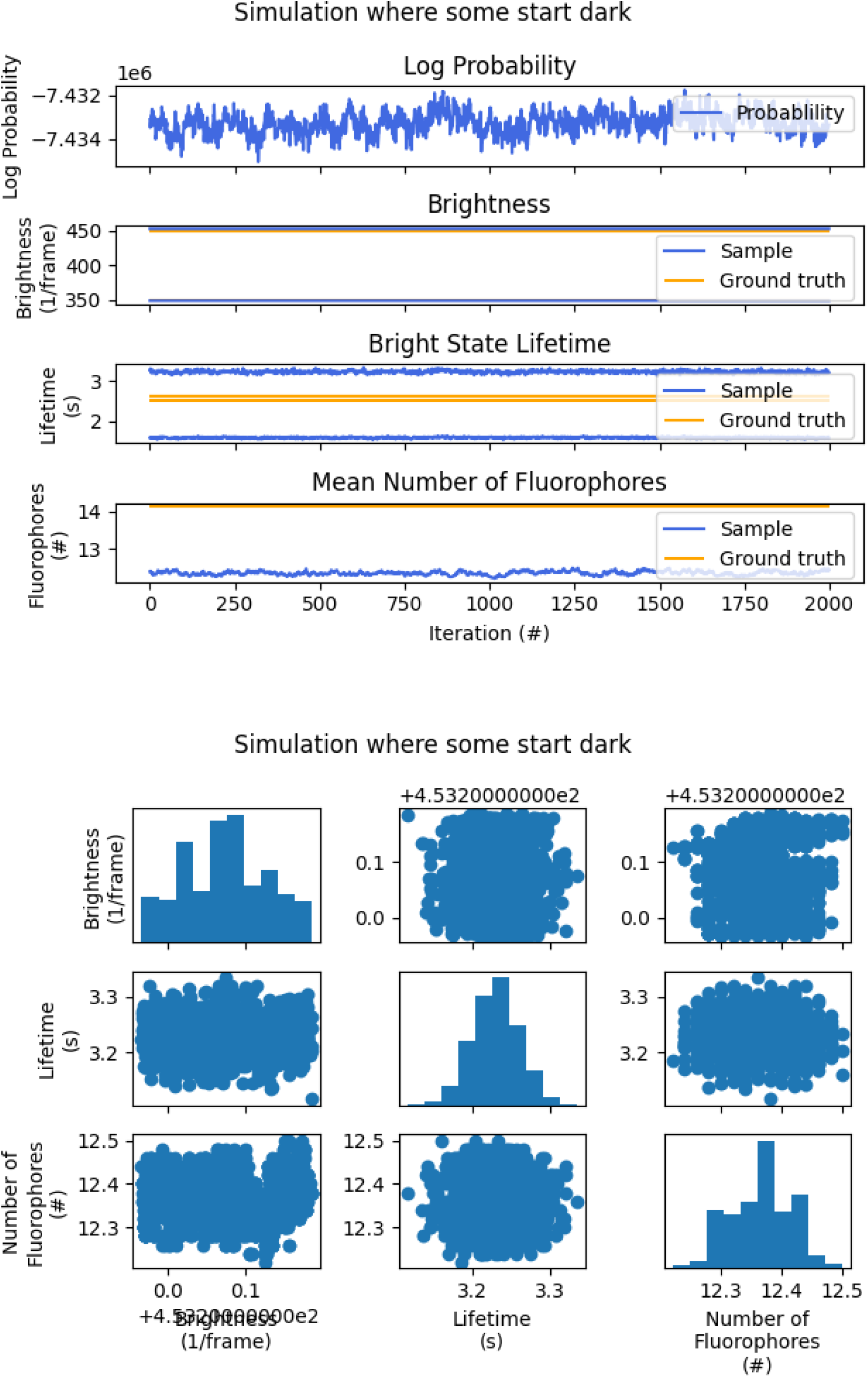

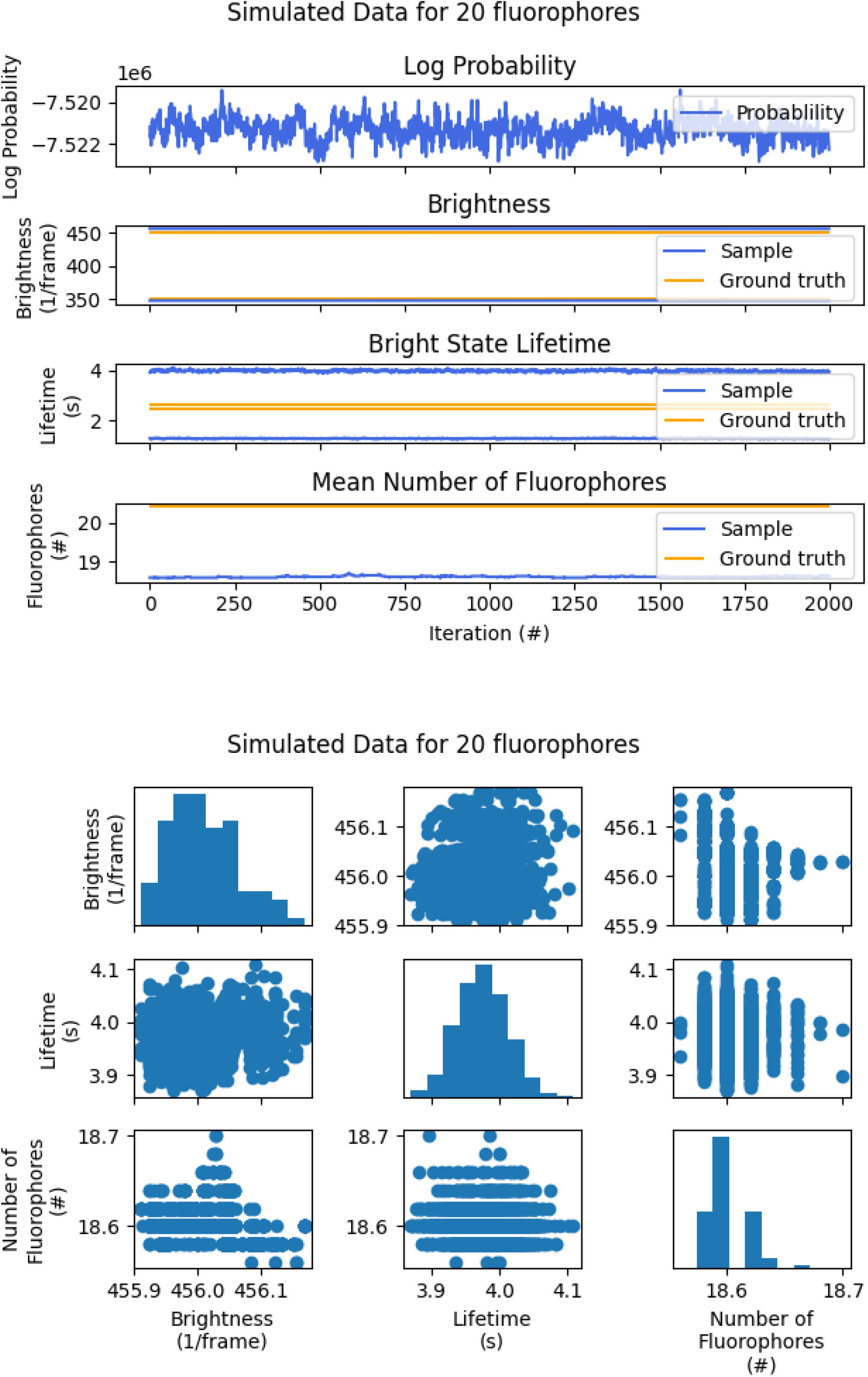

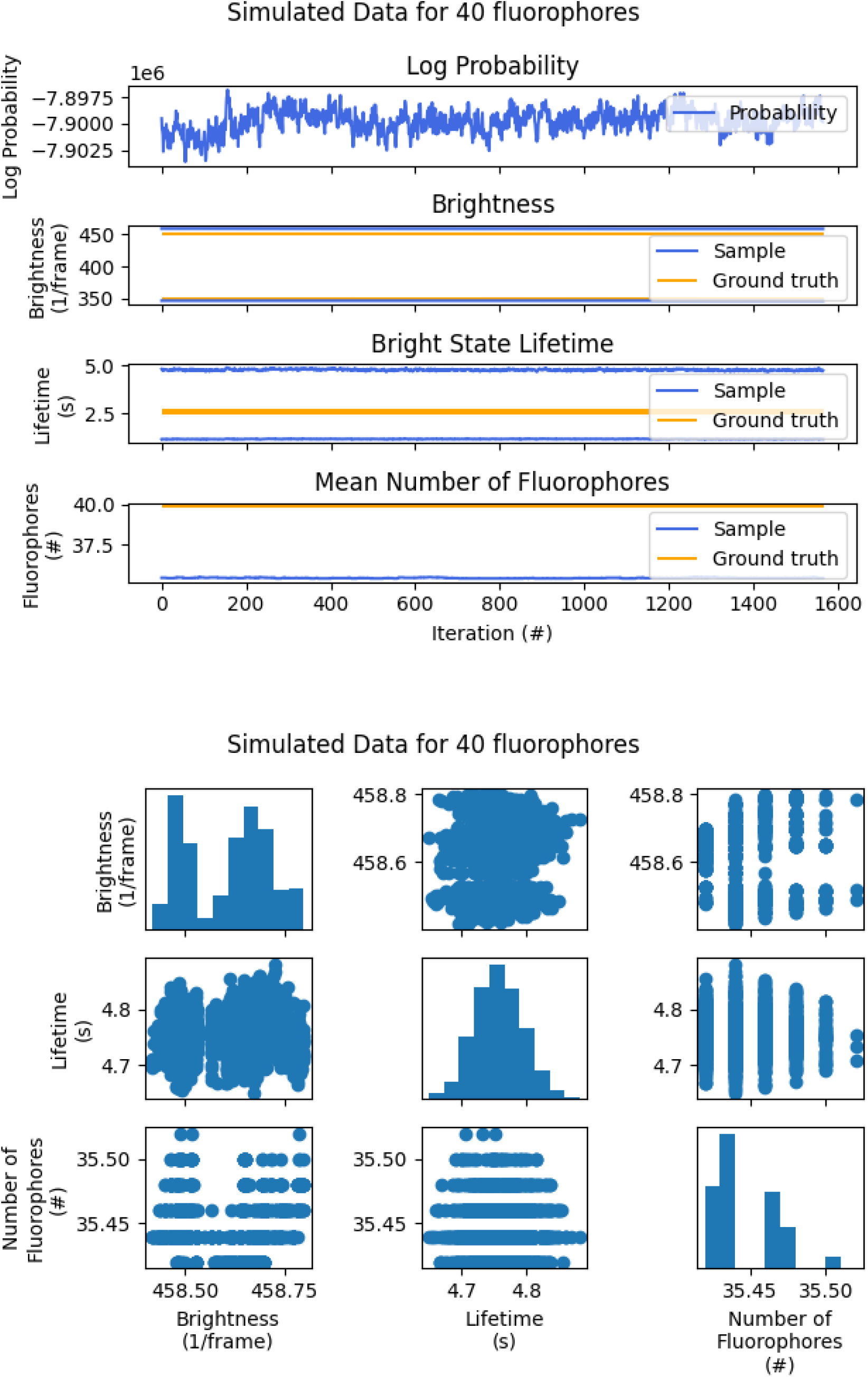

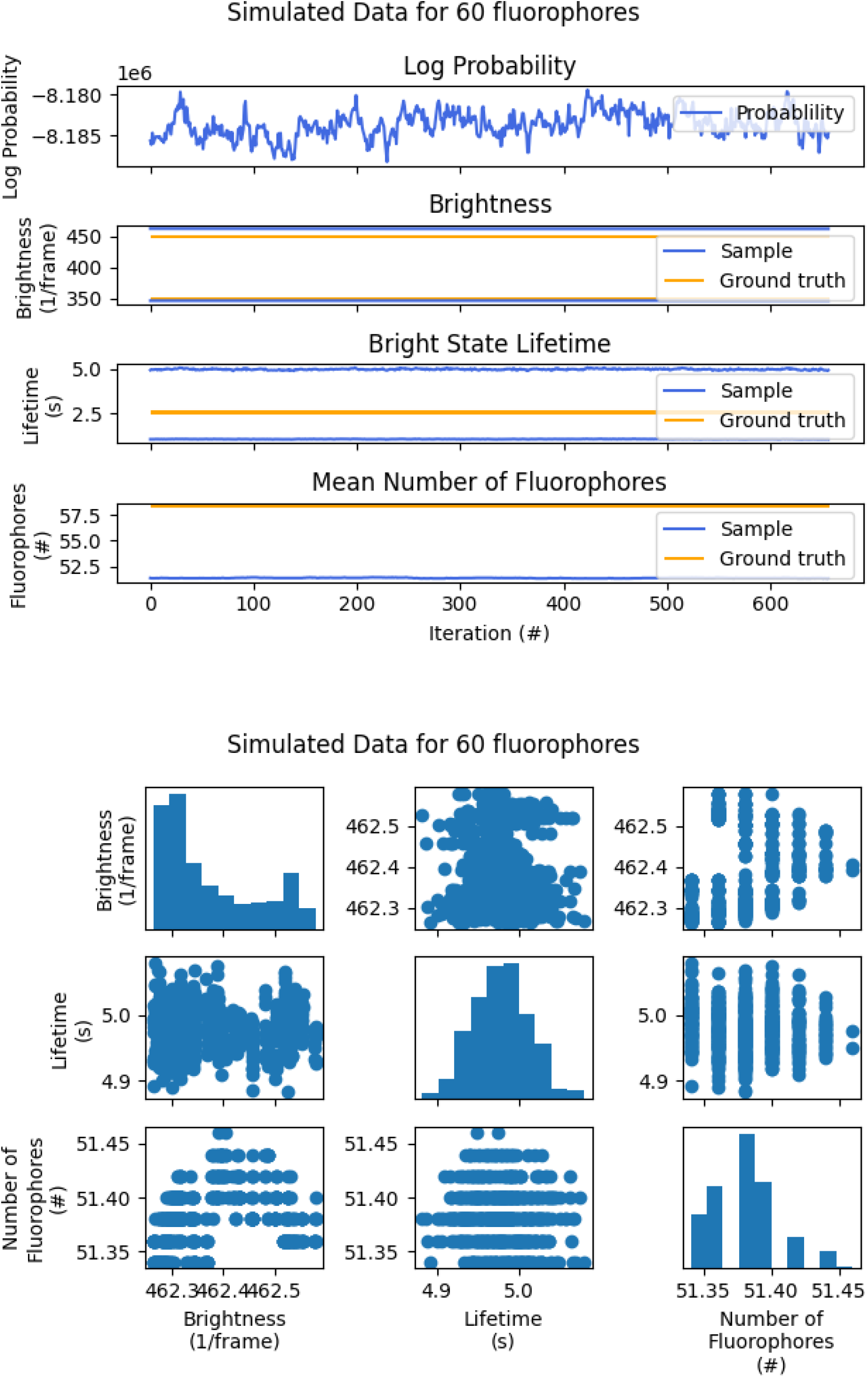

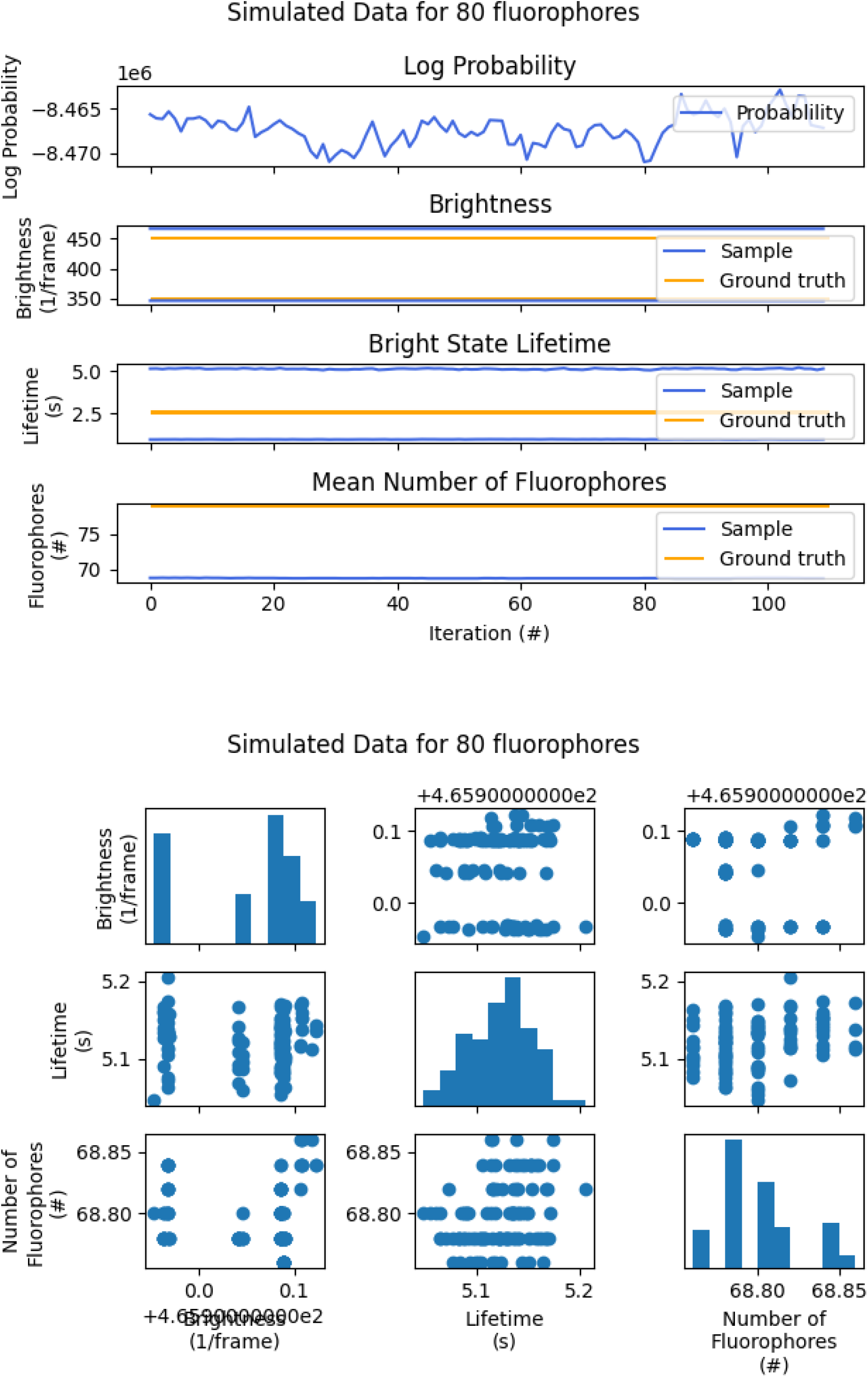

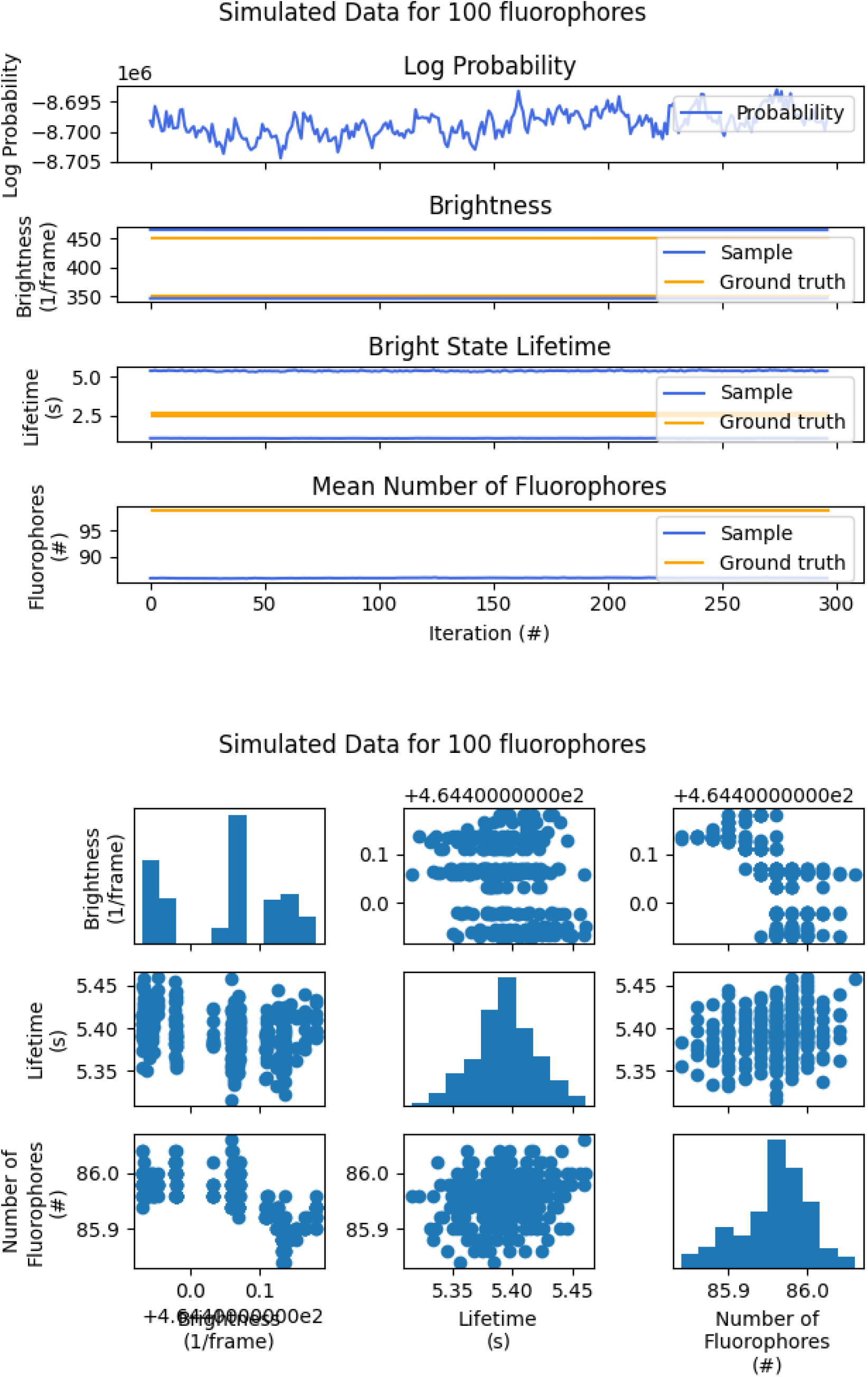

### 5.13 Plots of data

Here we plot the brightness vs time traces that we analyze in the main manuscript. Note that we only plot data from the first three ROIs, but there are, in fact, many more traces that we do not show. On the top under the title of each figure we print the total number of ROIs per data set.

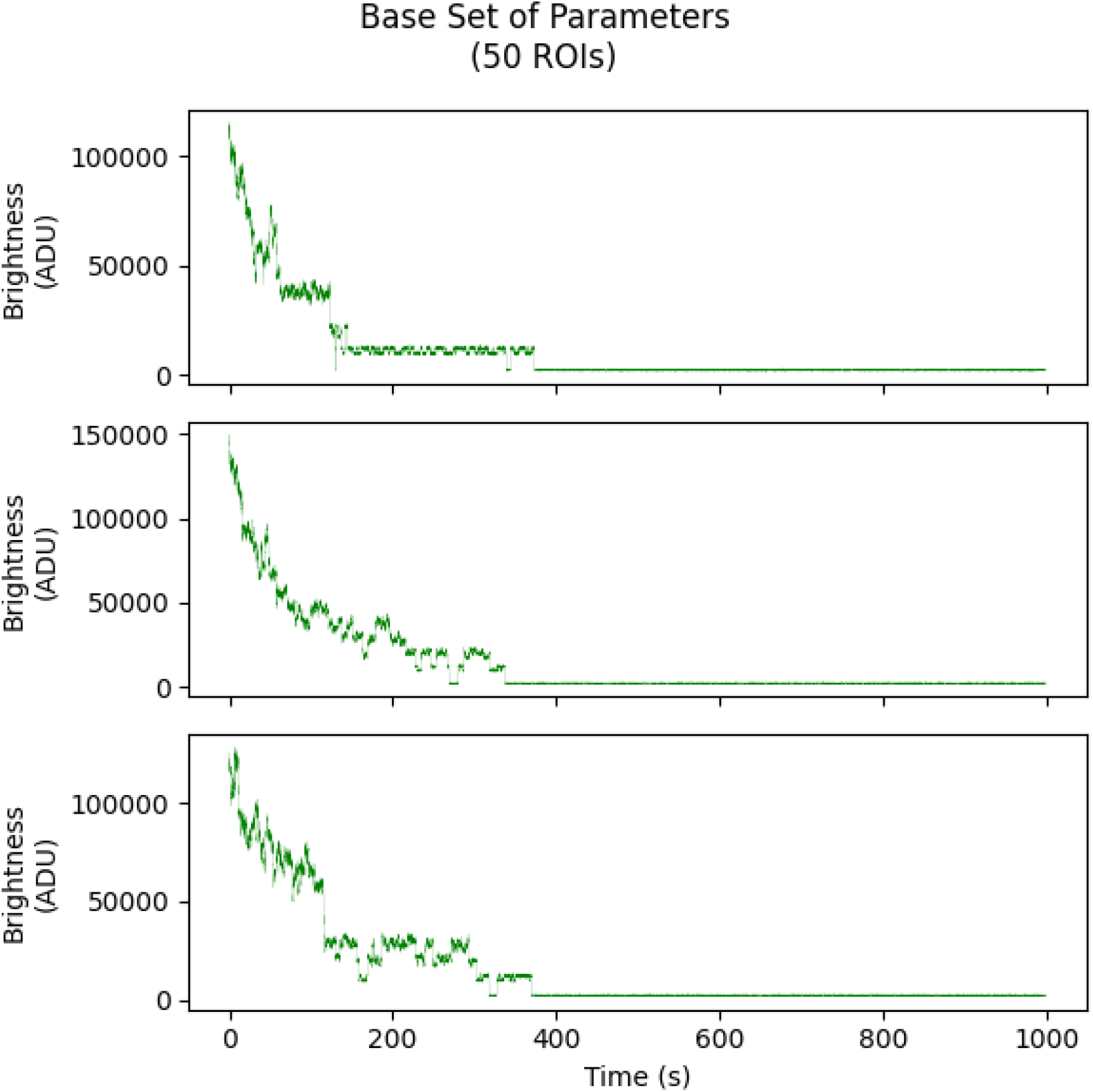

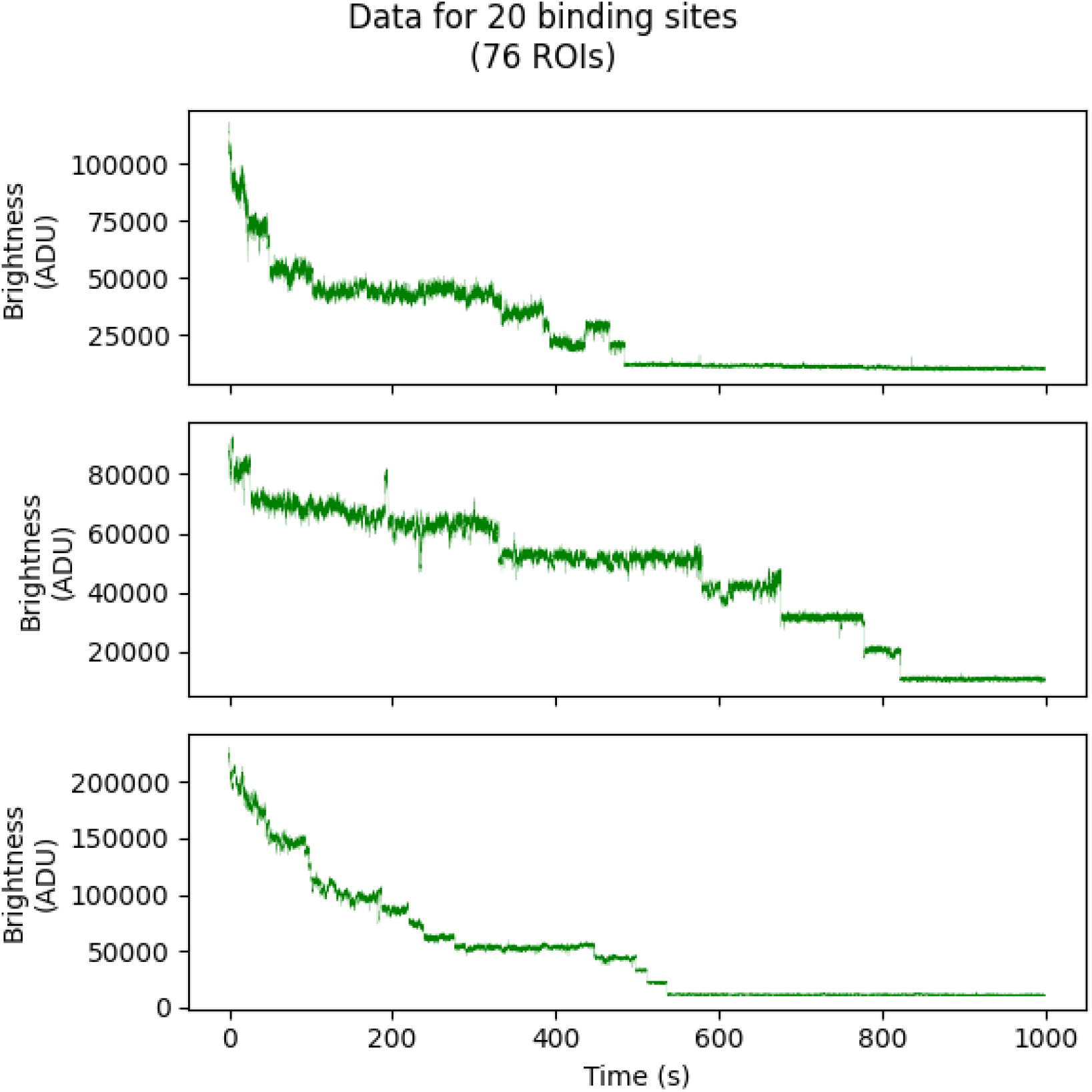

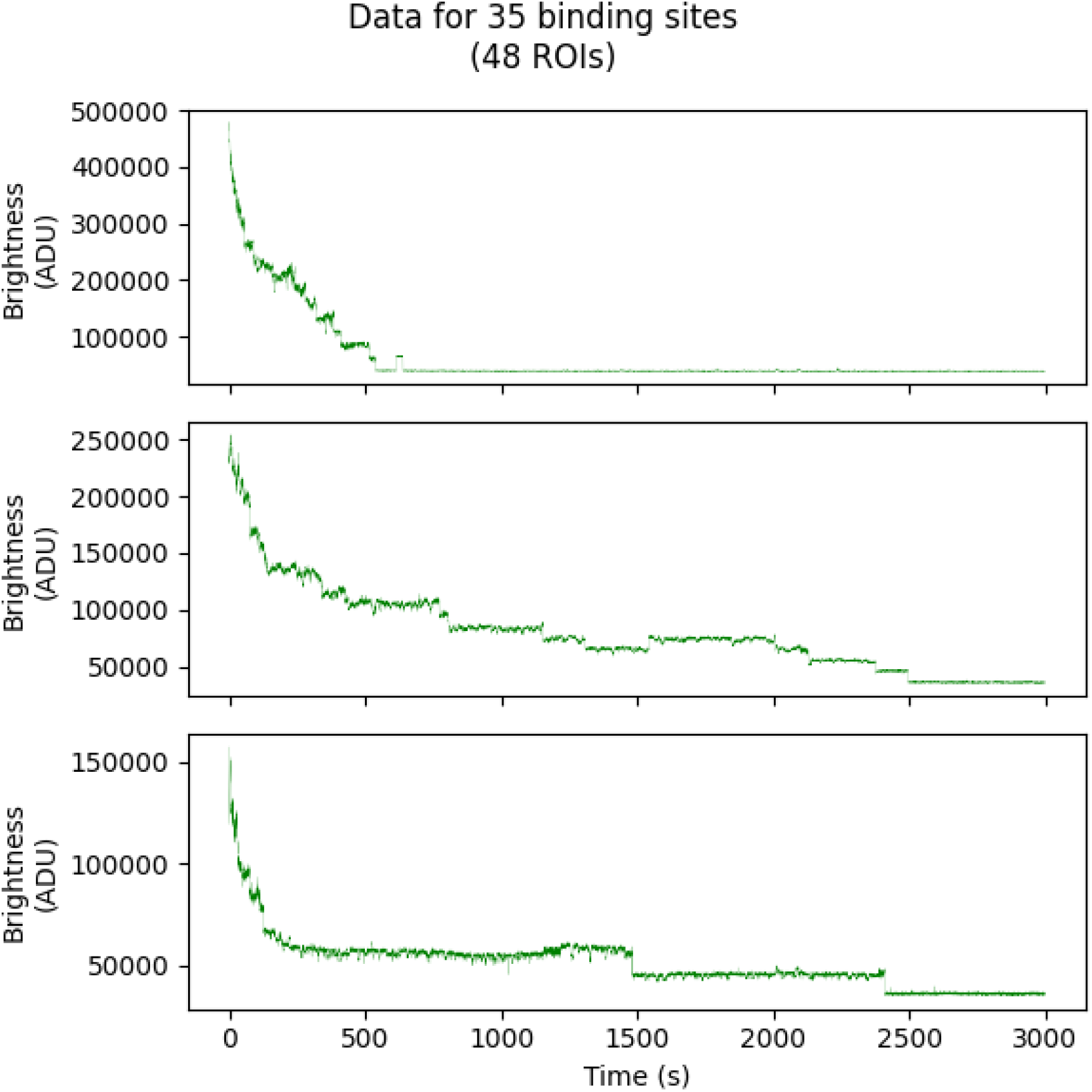

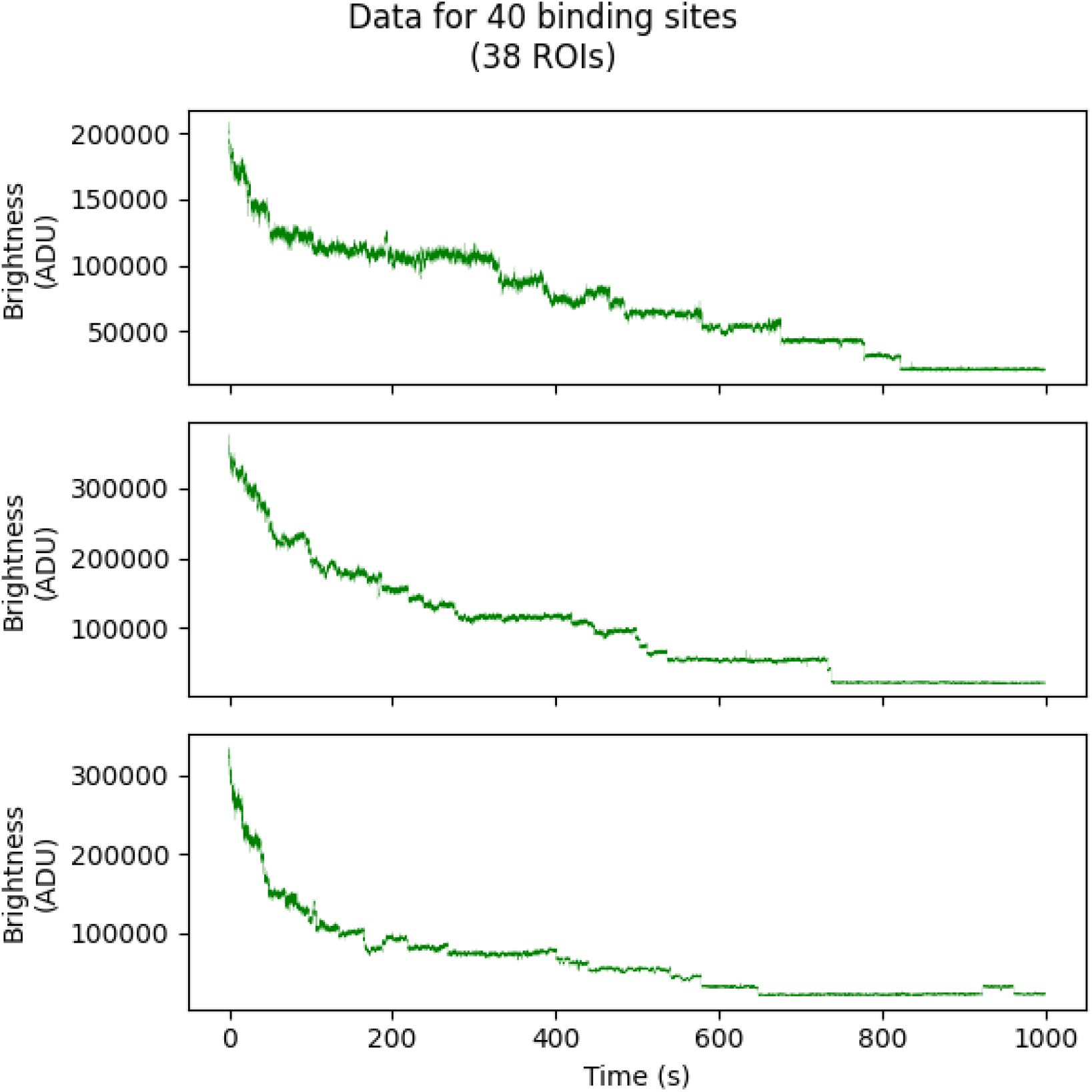

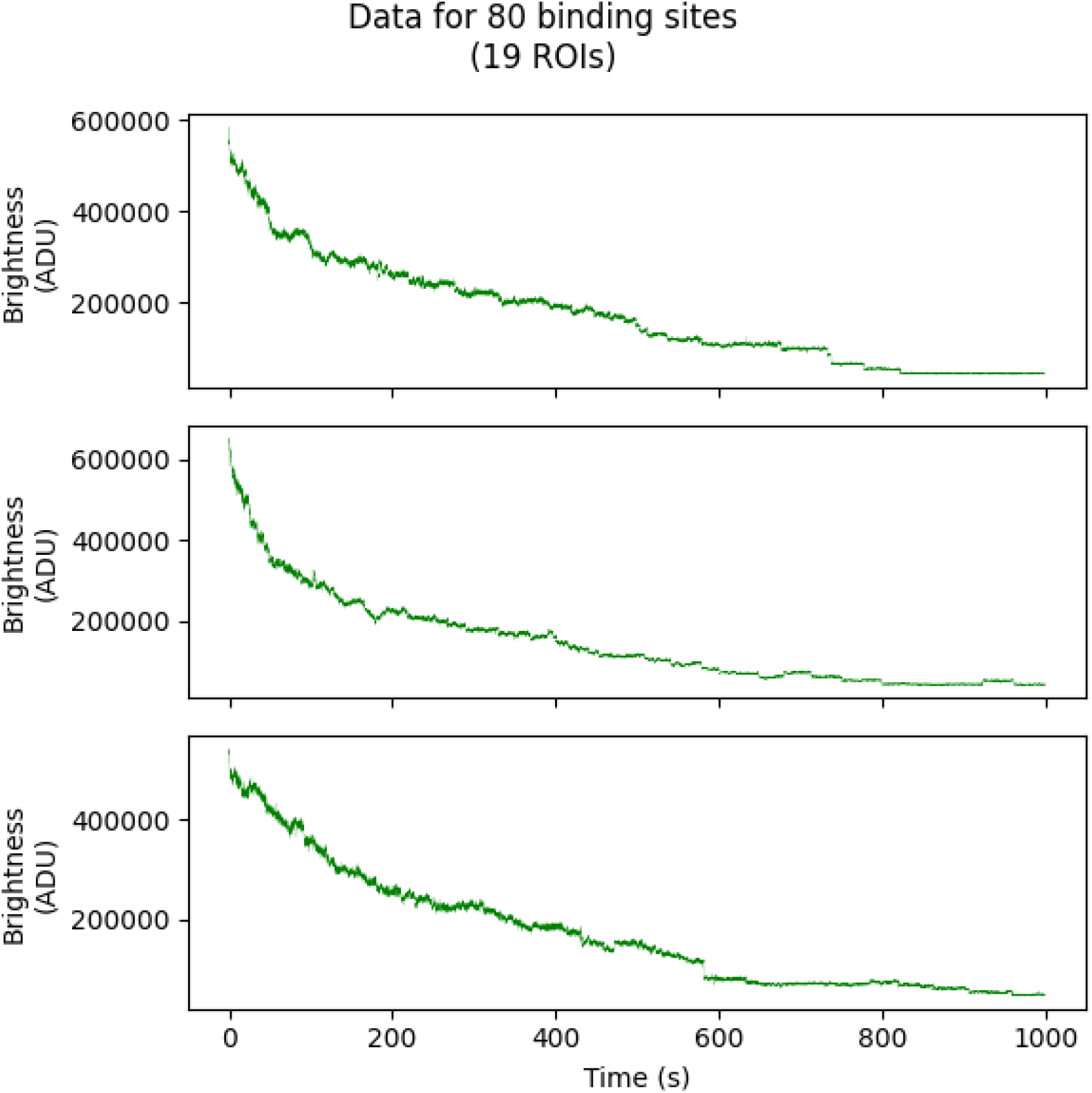

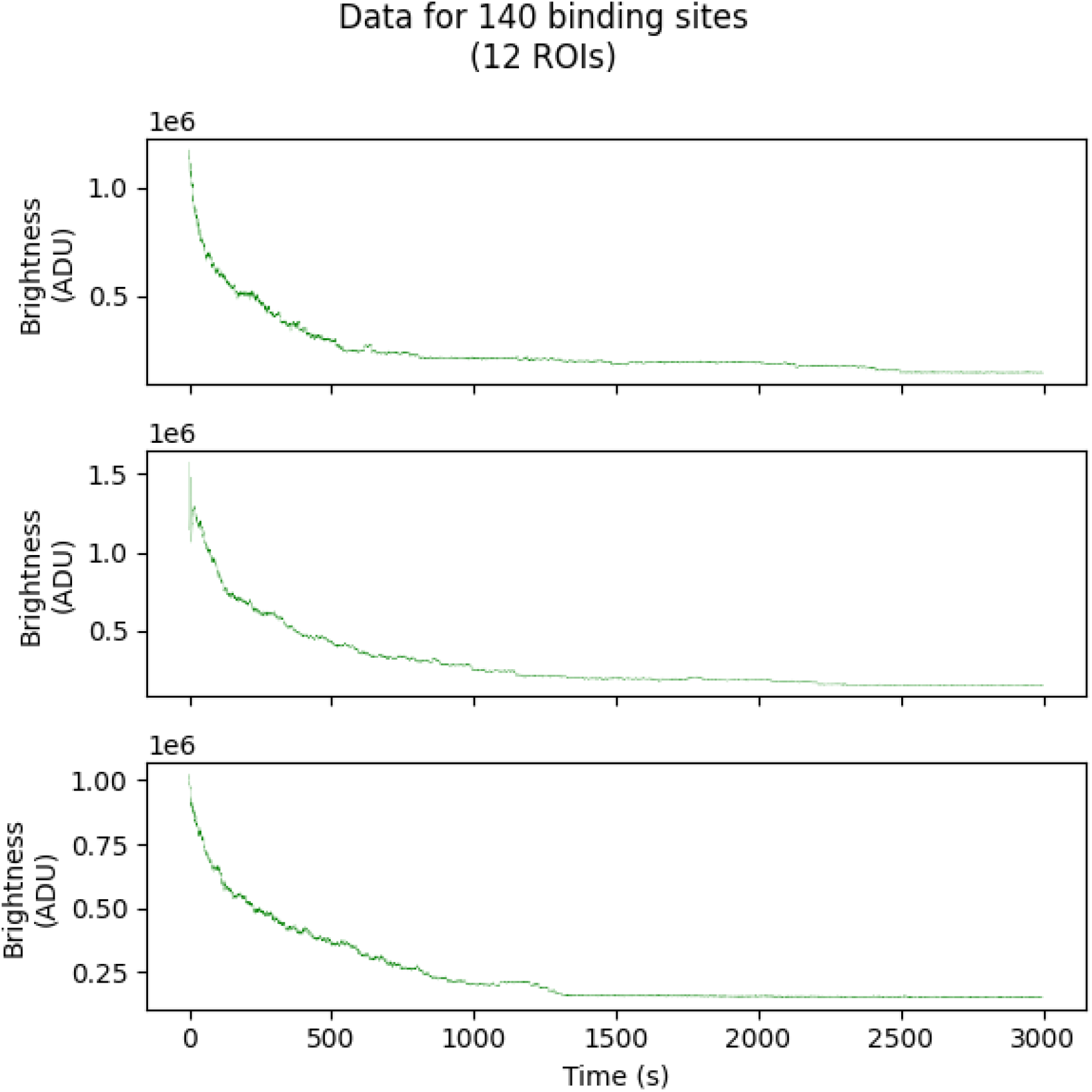

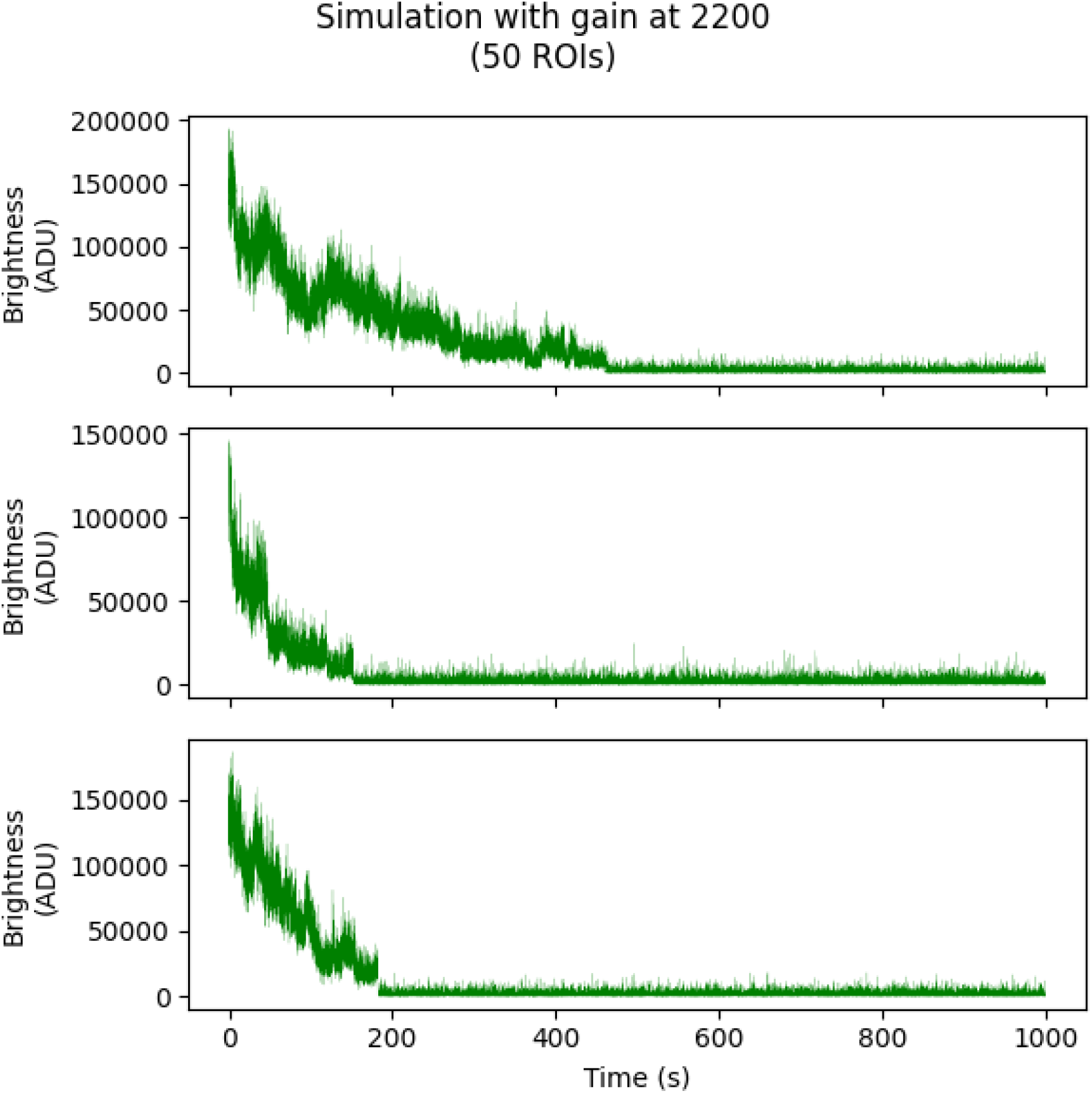

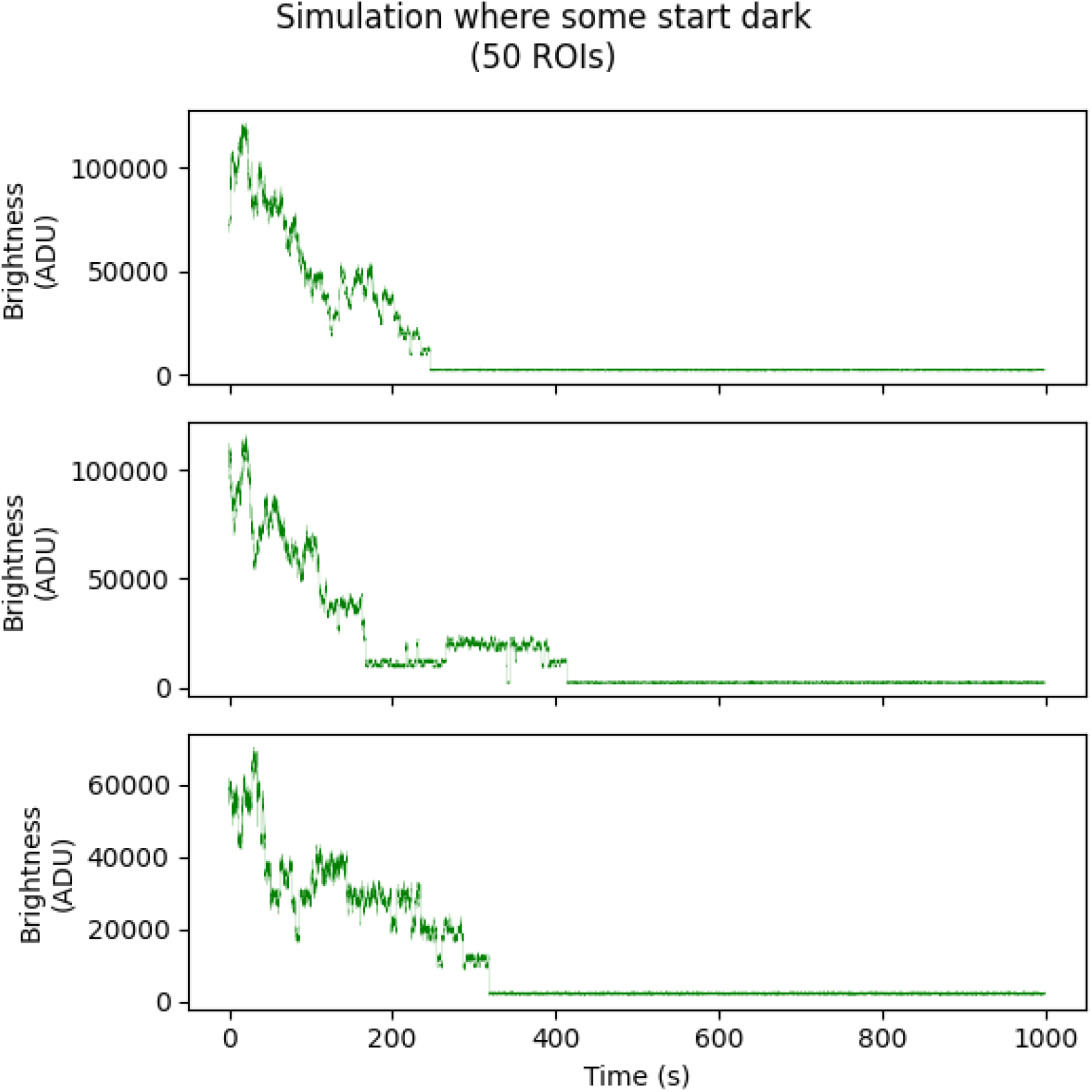

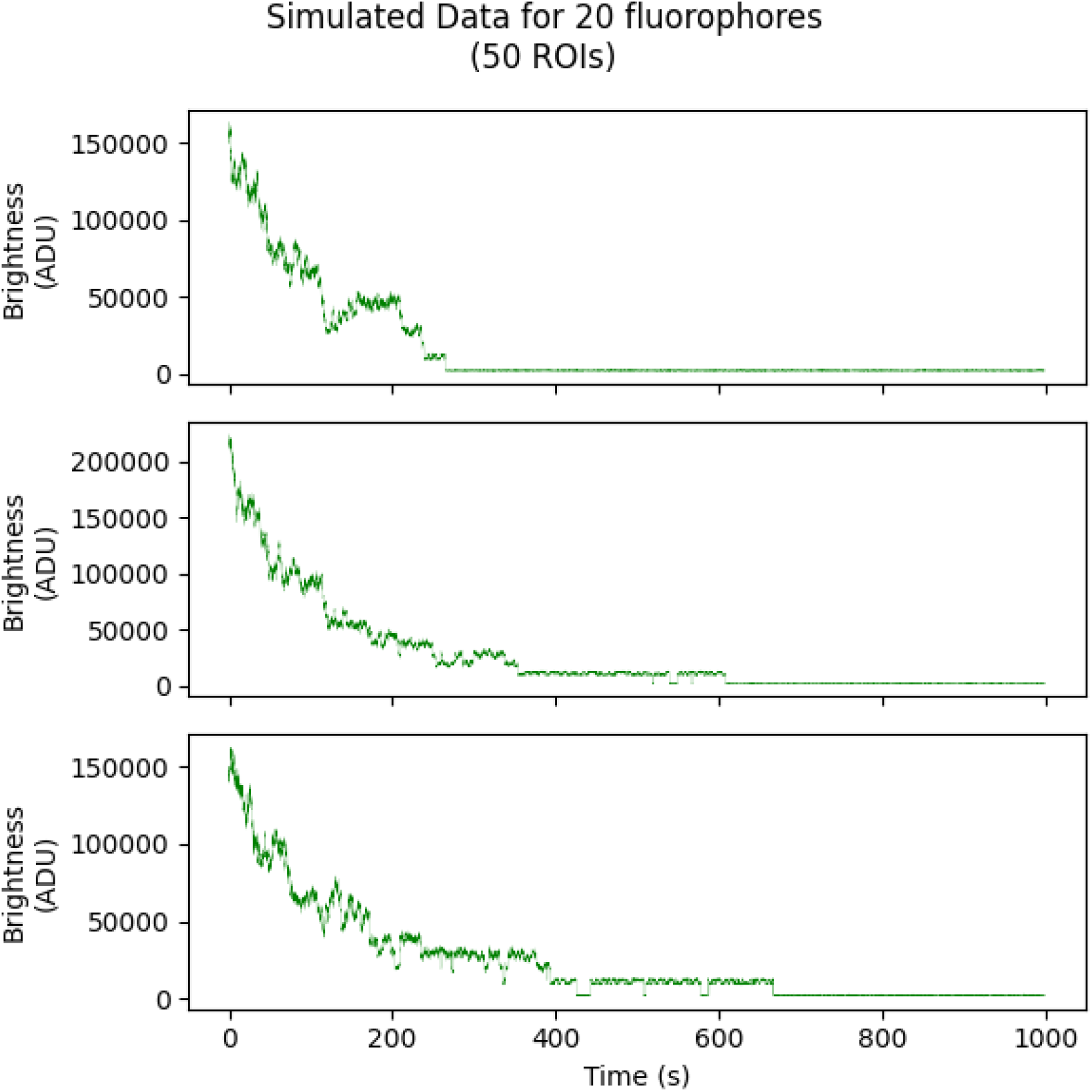

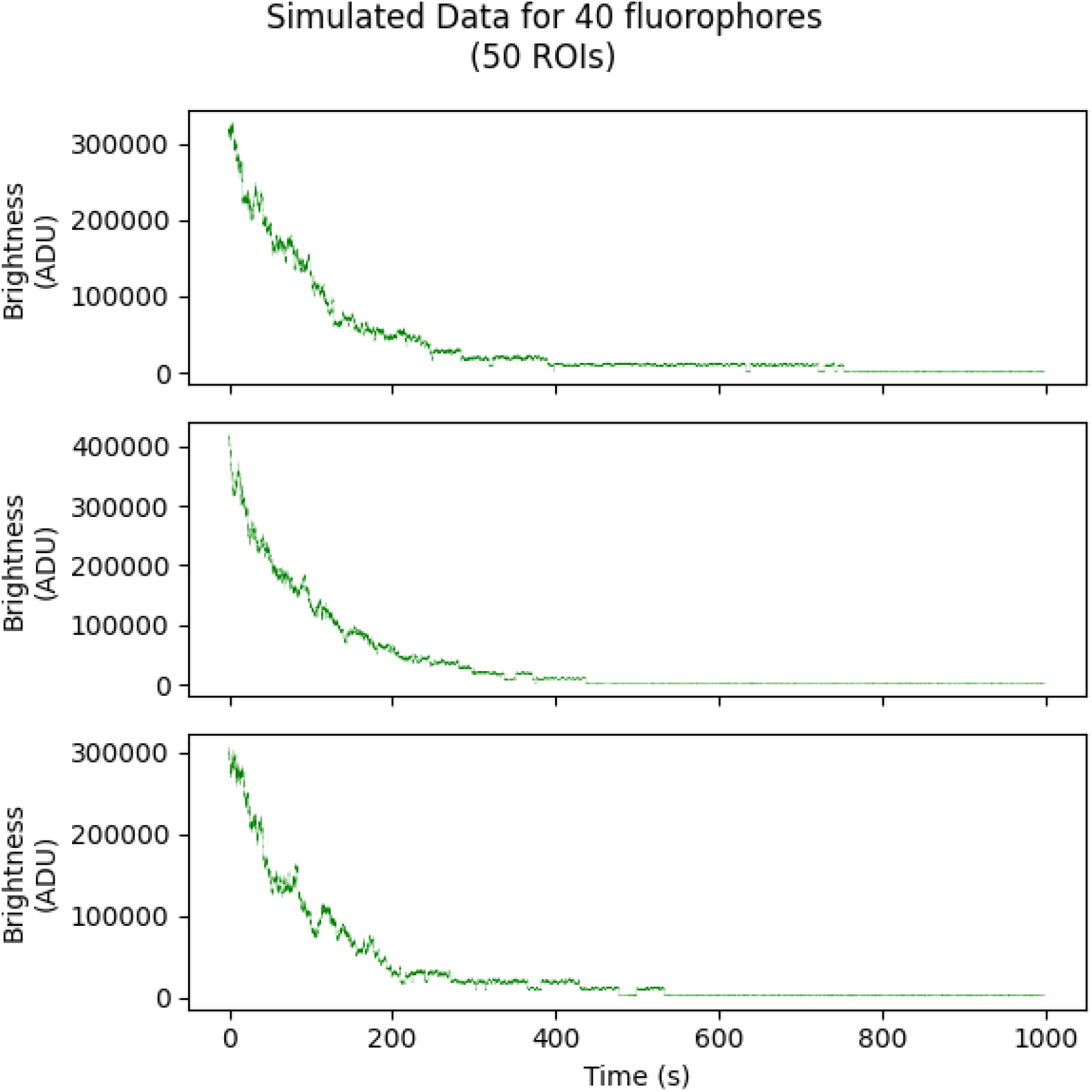

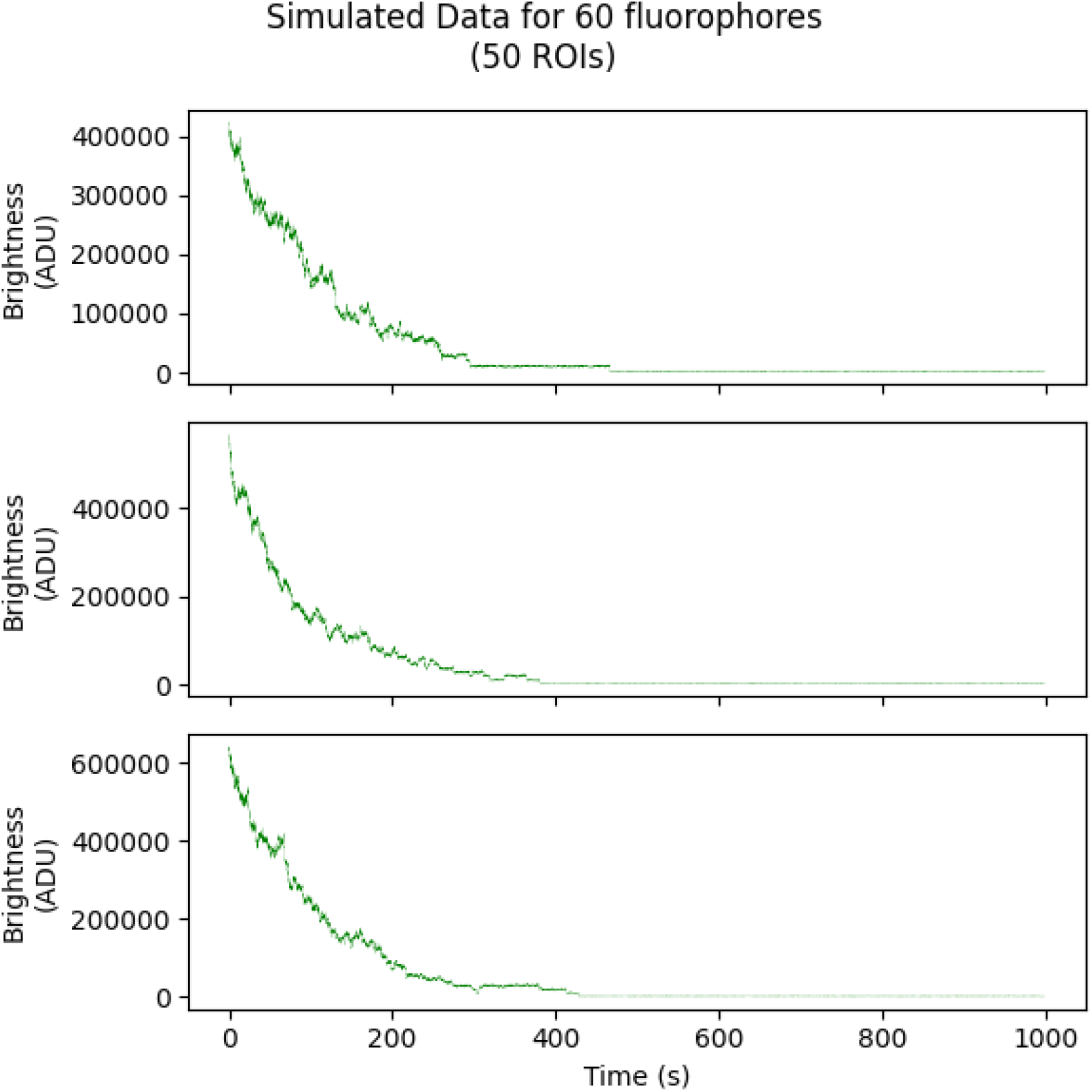

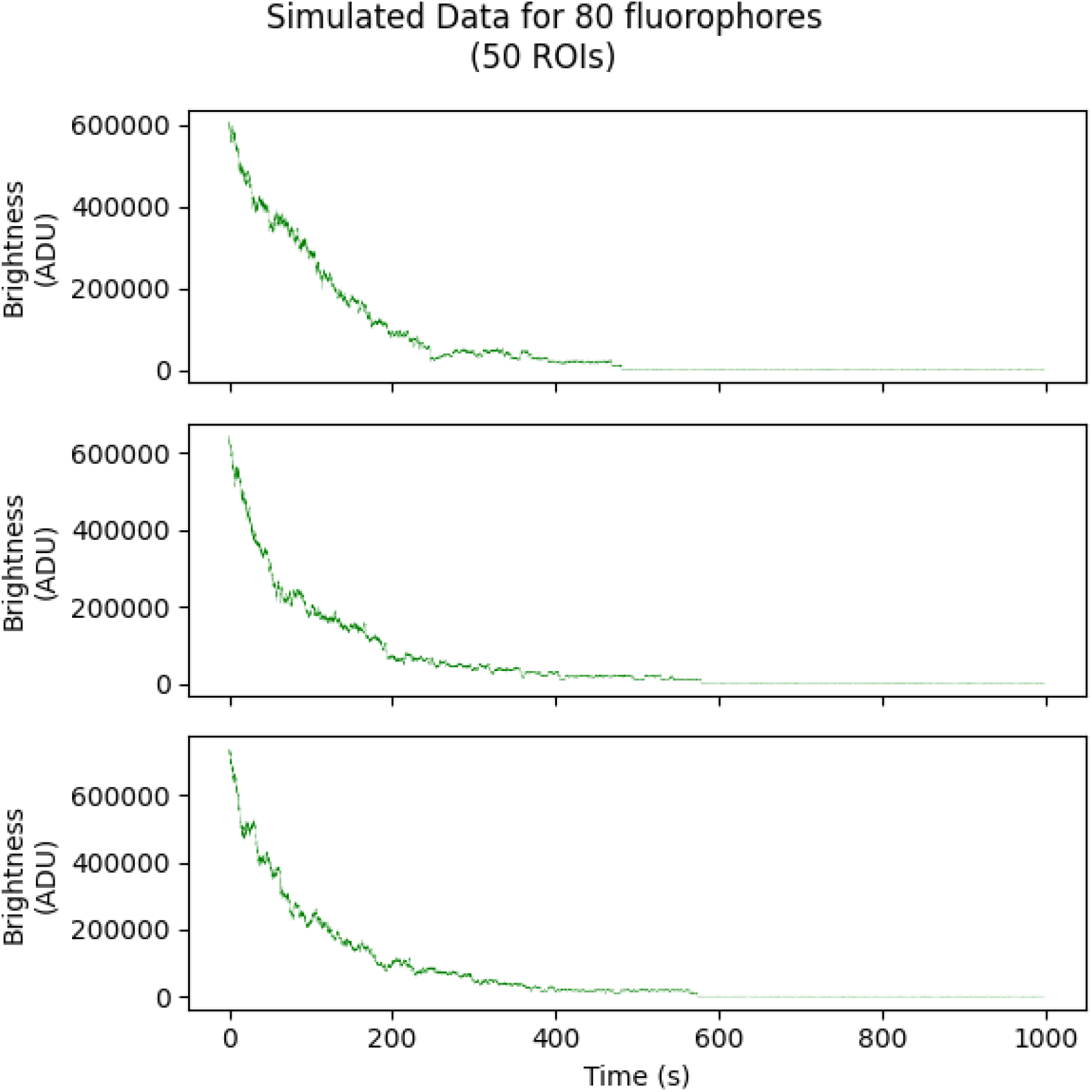

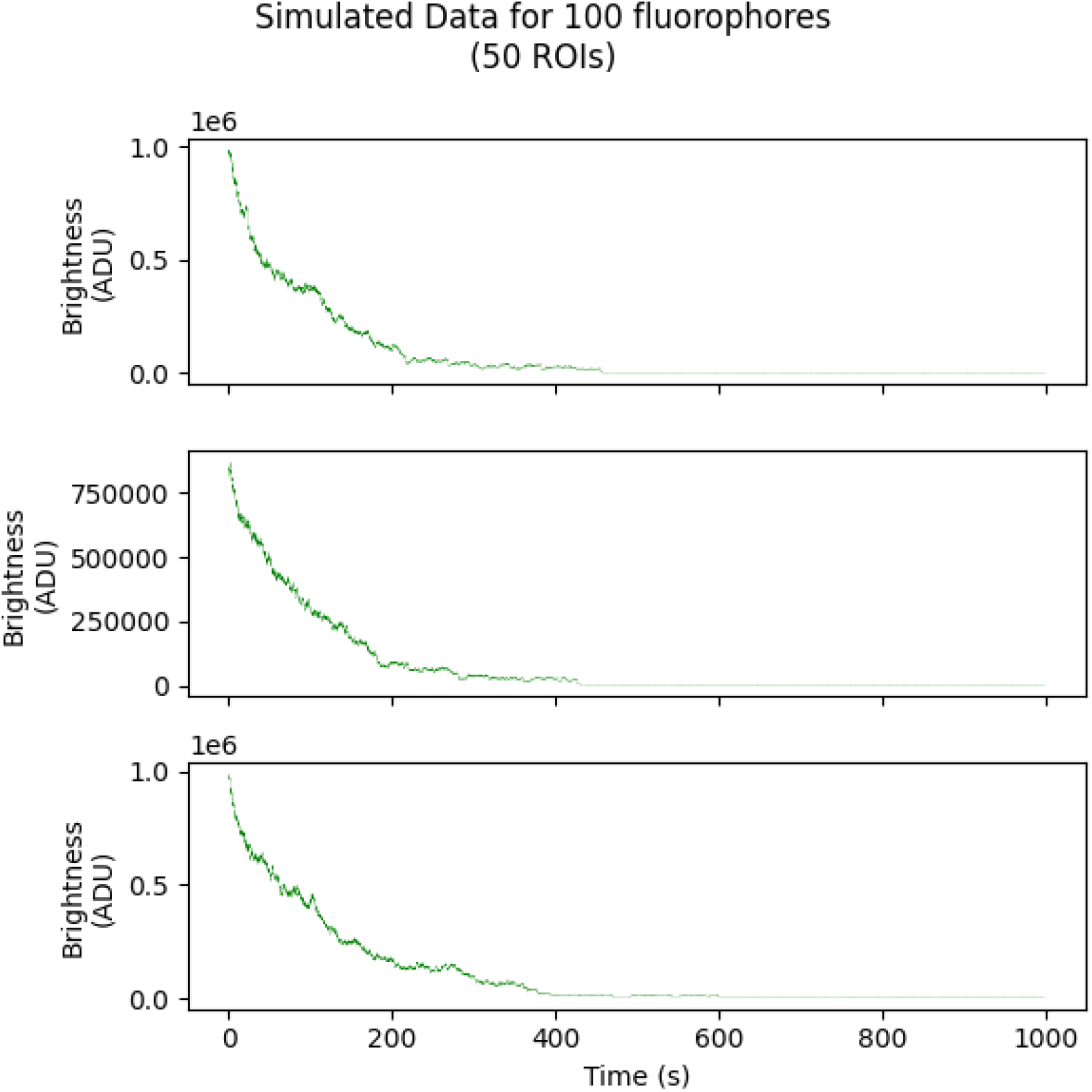

## References

[1] Seung-Ryoung Jung, Bryant S Fujimoto, and Daniel T Chiu. Quantitative microscopy based on single-molecule fluorescence. Current opinion in chemical biology, 39:64–73, 2017.

[2] Antony Lee, Konstantinos Tsekouras, Christopher Calderon, Carlos Bustamante, and Steve Pressé. Unraveling the thousand word picture: an introduction to super-resolution data analysis. Chemical reviews, 117(11):7276–7330, 2017.

[3] Bonnie O Leung and Keng C Chou. Review of super-resolution fluorescence microscopy for biology. Applied spectroscopy, 65(9):967–980, 2011.

[4] Won-Ki Cho, Jan-Hendrik Spille, Micca Hecht, Choongman Lee, Charles Li, Valentin Grube, and Ibrahim I Cisse. Mediator and RNA polymerase ii clusters associate in transcription-dependent condensates. Science, 361(6400):412–415, 2018.

[5] Ibrahim I Cisse, Ignacio Izeddin, Sebastien Z Causse, Lydia Boudarene, Adrien Senecal, Leila Muresan, Claire Dugast-Darzacq, Bassam Hajj, Maxime Dahan, and Xavier Darzacq. Real-time dynamics of RNA polymerase II clustering in live human cells. Science, 341(6146):664–667, 2013.

[6] Jervis Vermal Thevathasan, Maurice Kahnwald, Konstanty Cieśliński, Philipp Hoess, Sudheer Kumar Peneti, Manuel Reitberger, Daniel Heid, Krishna Chaitanya Kasuba, Sarah Janice Hoerner, Yiming Li, et al. Nuclear pores as versatile reference standards for quantitative superresolution microscopy. Nature methods, 16(10):1045–1053, 2019.

[7] Amanda Remorino, Simon De Beco, Fanny Cayrac, Fahima Di Federico, Gaetan Cornilleau, Alexis Gautreau, Maria Carla Parrini, Jean-Baptiste Masson, Maxime Dahan, and Mathieu Coppey. Gradients of rac1 nanoclusters support spatial patterns of rac1 signaling. Cell reports, 21(7):1922–1935, 2017.

[8] H Wang, X Yan, H Aigner, Andreas Bracher, Nghiem Dinh Nguyen, Wei Yih Hee, BM Long, Graeme Dean Price, FU Hartl, and M Hayer-Hartl. Rubisco condensate formation by ccmm in *β*-carboxysome biogenesis. Nature, 566(7742):131–135, 2019.

[9] Yang Eric Guo, John C Manteiga, Jonathan E Henninger, Benjamin R Sabari, Alessandra Dall’Agnese, Nancy M Hannett, Jan-Hendrik Spille, Lena K Afeyan, Alicia V Zamudio, Krishna Shrinivas, et al. Pol ii phosphorylation regulates a switch between transcriptional and splicing condensates. Nature, 572(7770):543–548, 2019.

[10] Salman F Banani, Hyun O Lee, Anthony A Hyman, and Michael K Rosen. Biomolecular condensates: organizers of cellular biochemistry. Nature reviews Molecular cell biology, 18(5):285–298, 2017.

[11] Mariano I Gabitto, Herve Marie-Nelly, Ari Pakman, Andras Pataki, Xavier Darzacq, and Michael I Jordan. A Bayesian nonparametric approach to super-resolution single-molecule localization. bioRxiv, 2020.

[12] Jolien Suzanne Verdaasdonk, Josh Lawrimore, and Kerry Bloom. Determining absolute protein numbers by quantitative fluorescence microscopy. In Methods in cell biology, volume 123, pages 347–365. Elsevier, 2014.

[13] Maximilian H Ulbrich and Ehud Y Isacoff. Subunit counting in membrane-bound proteins. Nature methods, 4(4):319, 2007.

[14] Timo Aspelmeier, Alexander Egner, and Axel Munk. Modern statistical challenges in high-resolution fluorescence microscopy. Annual Review of Statistics and Its Application, 2:163–202, 2015.

[15] Konstantinos Tsekouras, Thomas C Custer, Hossein Jashnsaz, Nils G Walter, and Steve Pressé. A novel method to accurately locate and count large numbers of steps by photobleaching. Molecular biology of the cell, 27(22):3601–3615, 2016.

[16] Keegan E Hines. Inferring subunit stoichiometry from single molecule photobleaching. The Journal of general physiology, 141(6):737–746, 2013.

[17] Hugo McGuire, Mark RP Aurousseau, Derek Bowie, and Rikard Blunck. Automating single subunit counting of membrane proteins in mammalian cells. Journal of Biological Chemistry, 287(43):35912–35921, 2012.

[18] Hui Zhang and Peixuan Guo. Single molecule photobleaching (smpb) technology for counting of rna, dna, protein and other molecules in nanoparticles and biological complexes by tirf instrumentation. Methods, 67(2):169–176, 2014.

[19] Bennett Kalafut and Koen Visscher. An objective, model-independent method for detection of nonuniform steps in noisy signals. Computer Physics Communications, 179(10):716–723, 2008.

[20] Yalei Chen, Nathan C Deffenbaugh, Charles T Anderson, and William O Hancock. Molecular counting by photobleaching in protein complexes with many subunits: best practices and application to the cellulose synthesis complex. Molecular biology of the cell, 25(22):3630–3642, 2014.

[21] Daniel Nino, Daniel Djayakarsana, and Joshua N Milstein. Nanoscopic stoichiometry and singlemolecule counting. Small Methods, page 1900082, 2019.

[22] Jiachao Xu, Gege Qin, Fang Luo, Lina Wang, Rong Zhao, Nan Li, Jinghe Yuan, and Xiaohong Fang. Automated stoichiometry analysis of single-molecule fluorescence imaging traces via deep learning. Journal of the American Chemical Society, 141(17):6976–6985, 2019.

[23] Thomas Staudt, Timo Aspelmeier, Oskar Laitenberger, Claudia Geisler, Alexander Egner, and Axel Munk. Statistical molecule counting in super-resolution fluorescence microscopy: Towards quantitative nanoscopy. arXiv preprint arXiv:1903.11577, 2019.

[24] Clarissa Liesche, Kristin S Grußmayer, Michael Ludwig, Stefan Wörz, Karl Rohr, Dirk-Peter Herten, Joël Beaudouin, and Roland Eils. Automated analysis of single-molecule photobleaching data by statistical modeling of spot populations. Biophysical journal, 109(11):2352–2362, 2015.

[25] Jon Garry, Yuchong Li, Brandon Shew, Claudiu C Gradinaru, and Andrew D Rutenberg. Bayesian counting of photobleaching steps with physical priors. The Journal of Chemical Physics, 152(2):024110, 2020.

[26] Johan Hummert, Klaus Yserentant, Theresa Fink, Jonas Euchner, and Dirk-Peter Herten. Photobleaching step analysis for robust determination of protein complex stoichiometries. bioRxiv, 2020.

[27] Valerie C Coffman and Jian-Qiu Wu. Counting protein molecules using quantitative fluorescence microscopy. Trends in biochemical sciences, 37(11):499–506, 2012.

[28] Xingbo Shi, Suli Dong, Minmin Li, Xiaojun Liu, Qingquan Zhang, Wenfeng Zhao, Chenghua Zong, Yewang Zhang, and Hongwei Gai. Counting quantum dot aggregates for the detection of biotinylated proteins. Chemical Communications, 51(12):2353–2356, 2015.

[29] Filip Delport, Ania Deres, Jun-ichi Hotta, Jeroen Pollet, Bert Verbruggen, Bert Sels, Johan Hofkens, and Jeroen Lammertyn. Improved method for counting dna molecules on biofunctionalized nanoparticles. Langmuir, 26(3):1594–1597, 2010.

[30] Kristina M Herbert, Susanta K Sarkar, Maria Mills, Hilda C Delgado De la Herran, Keir C Neuman, and Joan A Steitz. A heterotrimer model of the complete microprocessor complex revealed by singlemolecule subunit counting. Rna, 22(2):175–183, 2016.

[31] Paul D Simonson, Hannah A DeBerg, Pinghua Ge, John K Alexander, Okunola Jeyifous, William N Green, and Paul R Selvin. Counting bungarotoxin binding sites of nicotinic acetylcholine receptors in mammalian cells with high signal/noise ratios. Biophysical journal, 99(10):L81–L83, 2010.

[32] Nela Durisic, Antoine G Godin, Claudia M Wever, Colin D Heyes, Melike Lakadamyali, and Joseph A Dent. Stoichiometry of the human glycine receptor revealed by direct subunit counting. Journal of Neuroscience, 32(37):12915–12920, 2012.

[33] Adam D Cawte, Peter J Unrau, and David S Rueda. Live cell imaging of single RNA molecules with fluorogenic mango ii arrays. bioRxiv, page 840082, 2019.

[34] Parul Agrawal, Anthony L DeVico, James S Foulke Jr, George K Lewis, Marzena Pazgier, and Krishanu Ray. Stoichiometric analyses of soluble cd4 to native-like hiv-1 envelope by single-molecule fluorescence spectroscopy. Cell Reports, 29(1):176–186, 2019.

[35] Antony Ann-Tzer Lee. Improving the resolution and accuracy of optical tweezers through algorithmic and instrumental advances. PhD thesis, UC Berkeley, 2018.

[36] Yuchong Li. Insane in the Membrane: The Functional Assembly of a G Protein Coupled Receptor at the Single-Molecule Level. PhD thesis, University of Toronto, 2018.

[37] Ved Prakash, Konstantinos Tsekouras, Muthukumaran Venkatachalapathy, Laurie Heinicke, Steve Pressé, Nils G Walter, and Yamuna Krishnan. Quantitative mapping of endosomal dna processing by single molecule counting. Angewandte Chemie International Edition, 58(10):3073–3076, 2019.

[38] Marijonas Tutkus, Parveen Akhtar, Jevgenij Chmeliov, Fanni Gorfol, Gediminas Trinkunas, Petar H Lambrev, and Leonas Valkunas. Fluorescence microscopy of single liposomes with incorporated pigment–proteins. Langmuir, 34(47):14410–14418, 2018.

[39] Amani A Hariri, Graham D Hamblin, Jack S Hardwick, Robert Godin, Jean-Francois Desjardins, Paul W Wiseman, Hanadi F Sleiman, and Gonzalo Cosa. Stoichiometry and dispersity of dna nanostructures using photobleaching pair-correlation analysis. Bioconjugate chemistry, 28(9):2340–2349, 2017.

[40] Andrew Gelman, John B Carlin, Hal S Stern, David B Dunson, Aki Vehtari, and Donald B Rubin. Bayesian data analysis. Chapman and Hall/CRC, 2013.

[41] Udo Von Toussaint. Bayesian inference in physics. Reviews of Modern Physics, 83(3):943, 2011.

[42] Mingyuan Zhou, Lauren Hannah, David Dunson, and Lawrence Carin. Beta-negative binomial process and poisson factor analysis. In Artificial Intelligence and Statistics, pages 1462–1471, 2012.

[43] Romain Thibaux and Michael I Jordan. Hierarchical beta processes and the Indian buffet process. In Artificial Intelligence and Statistics, pages 564–571, 2007.

[44] Michael Hirsch, Richard J Wareham, Marisa L Martin-Fernandez, Michael P Hobson, and Daniel J Rolfe. A stochastic model for electron multiplication charge-coupled devices–from theory to practice. PloS one, 8(1):e53671, 2013.

[45] Fang Huang, Tobias MP Hartwich, Felix E Rivera-Molina, Yu Lin, Whitney C Duim, Jane J Long, Pradeep D Uchil, Jordan R Myers, Michelle A Baird, Walther Mothes, et al. Video-rate nanoscopy using scmos camera–specific single-molecule localization algorithms. Nature methods, 10(7):653, 2013.

[46] Mark Stanford Robbins and Benjamin James Hadwen. The noise performance of electron multiplying charge-coupled devices. IEEE transactions on electron devices, 50(5):1227–1232, 2003.

[47] Jurgen V Gael, Yee W Teh, and Zoubin Ghahramani. The infinite factorial hidden Markov model. In Advances in Neural Information Processing Systems, pages 1697–1704, 2009.

[48] Christopher M Bishop. Pattern recognition and machine learning. springer, 2006.

[49] Meysam Tavakoli, Sina Jazani, Ioannis Sgouralis, Omer M Shafraz, Sanjeevi Sivasankar, Bryan Donaphon, Marcia Levitus, and Steve Pressé. Pitching single-focus confocal data analysis one photon at a time with Bayesian nonparametrics. Physical Review X, 10(1):011021, 2020.

[50] Sina Jazani, Ioannis Sgouralis, Omer M Shafraz, Marcia Levitus, Sanjeevi Sivasankar, and Steve Pressé. An alternative framework for fluorescence correlation spectroscopy. Nature communications, 10(1):1–10, 2019.

[51] Walter R Gilks and Pascal Wild. Adaptive rejection sampling for gibbs sampling. Journal of the Royal Statistical Society: Series C (Applied Statistics), 41(2):337–348, 1992.

[52] Stuart Geman and Donald Geman. Stochastic relaxation, gibbs distributions, and the Bayesian restoration of images. IEEE Transactions on pattern analysis and machine intelligence, pages 721–741, 1984.

[53] Christian Robert and George Casella. Monte Carlo statistical methods. Springer Science & Business Media, 2013.

[54] Viktorija Glembockyte, Junan Lin, and Gonzalo Cosa. Improving the photostability of red-and green-emissive single-molecule fluorophores via ni2+ mediated excited triplet-state quenching. The Journal of Physical Chemistry B, 120(46):11923–11929, 2016.

[55] Huimin Lu, Hui Zhang, Shaowu Yang, and Zhiqiang Zheng. Camera parameters auto-adjusting technique for robust robot vision. In 2010 IEEE International Conference on Robotics and Automation, pages 1518–1523. IEEE, 2010.

[56] Didier Casanova, Domitille Giaume, Mélanie Moreau, Jean-Louis Martin, Thierry Gacoin, Jean-Pierre Boilot, and Antigoni Alexandrou. Counting the number of proteins coupled to single nanoparticles. Journal of the American Chemical Society, 129(42):12592–12593, 2007.

[57] Geoffrey C Rollins, Jae Yen Shin, Carlos Bustamante, and Steve Pressé. Stochastic approach to the molecular counting problem in superresolution microscopy. Proceedings of the National Academy of Sciences, 112(2):E110–E118, 2015.

[58] Hari Shroff, Helen White, and Eric Betzig. Photoactivated localization microscopy (palm) of adhesion complexes. Current protocols in cell biology, 41(1):4–21, 2008.

[59] Michael J Rust, Mark Bates, and Xiaowei Zhuang. Stochastic optical reconstruction microscopy (storm) provides sub-diffraction-limit image resolution. Nature methods, 3(10):793, 2006.

[60] John E Wampler and Karl Kutz. Quantitative fluorescence microscopy using photomultiplier tubes and imaging detectors. Methods in cell biology, 29:239–267, 1988.

[61] Zhaoning Zhang, Yujie Wang, Rafael Piestun, and Zhen-Li Huang. Characterizing and correcting camera noise in back-illuminated scmos cameras. Optics Express, 29(5):6668–6690, 2021.

[62] Ioannis Sgouralis and Steve Pressé. An introduction to infinite HMMs for single-molecule data analysis. Biophysical journal, 112(10):2021–2029, 2017.

[63] Ioannis Sgouralis and Steve Pressé. Icon: an adaptation of infinite HMMs for time traces with drift. Biophysical journal, 112(10):2117–2126, 2017.

[64] Zeliha Kilic, Ioannis Sgouralis, and Steve Presse. Residence time analysis of RNA polymerase transcription dynamics: A Bayesian sticky HMM approach. bioRxiv, 2020.

[65] Devinderjit Sivia and John Skilling. Data analysis: a Bayesian tutorial. OUP Oxford, 2006.

[66] Michael Betancourt. A conceptual introduction to Hamiltonian Monte Carlo. arXiv preprint arXiv:1701.02434, 2017.

